# Population structure and domestication history of the Javan banteng (*Bos javanicus javanicus*)

**DOI:** 10.1101/2025.04.01.646613

**Authors:** Xi Wang, Sabhrina Gita Aninta, Genís Garcia-Erill, Zilong Li, Anubhab Khan, Xiaodong Liu, Laura D. Bertola, Anik Budhi Dharmayanthi, Yulianto, Yonathan, Conor Rossi, Reagan Cauble-Sims, Benjamin D. Rosen, Darren E. Hagen, Michael P. Heaton, Timothy P. L. Smith, Johannes A. Lenstra, Nuno F.G. Martins, Mikkel-Holger S. Sinding, Muhammad Agil, Bambang Purwantara, Christina Hvilsom, Gono Semiadi, Rasmus Heller

## Abstract

The domestication of the banteng in Southeast Asia is one of the World’s least known livestock domestications, yet a vital component of the agricultural system in Indonesia and surrounding countries. Here we generated the first reference genome of the banteng and used it to analyze a set of 78 resequenced wild and domesticated bantengs, including 19 newly generated whole-genome sequenced samples of which three are historical samples. We found low heterozygosity and significant differentiation, the latter primarily driven by recent genetic drift and inbreeding in two populations, and clearly attributable to anthropogenically driven founder events or *ex-situ* breeding. Population structure when excluding these two populations was limited, and we found that the evolutionary divergence between wild and domestic banteng is moderate (*F*_ST_ = 0.14), relatively young (10,356 years), and with post-divergence gene flow. We found only weak signals of a domestication bottleneck between ∼6100-2900 years ago, and genetic diversity is on average higher in domestic than in wild banteng. Despite the soft domestication history, we found 56 candidate genes under selection during domestication, with the leptin receptor gene (*LEPR*) of particular interest due to the robust selection signal across methods, and its known association to metabolism, obesity and energy homeostasis. Finally, genetic load estimation revealed that Bali cattle in Australia have high realized load, while Bali cattle from Bali have high masked load. These findings provide the first genomic insights into an understudied bovine that is Critically Endangered in its wild form, and agriculturally important in its domesticated form.

## Introduction

Domestication of plants and animals is one of the most important developments in human history (Diamond 2002; Larson et al. 2014; Purugganan 2022). At the beginning of the Holocene, about 11,000 years before present (YBP), many human societies intensified their socioeconomic transition from hunting and gathering to the cultivation of plants and herding of animals, leading to the domestication of crops and livestock (Diamond 2002; Larson et al. 2014; Purugganan 2022). Investigating when, where, and how domestication of different species took place is therefore essential for understanding the origins of complex human civilizations (Diamond 2002; Larson et al. 2014; Purugganan 2022). Genetic changes and phenotypic diversification underlying domestication have also provided a key model for biologists to study evolutionary mechanisms (Frantz et al. 2020). Accordingly, most of the livestock species utilized by humans today have relatively well-characterized evolutionary histories, although there are notable exceptions.

The genus *Bos* consists of 8-9 species and experienced an unusually high prevalence of unique domestications with at least four independent domestication events (Wu et al. 2018). While the domestication history of cattle (*Bos taurus*, *Bos indicus*) and of yak (*B. mutus*) are well-known (Achilli et al. 2008; Qiu et al. 2015; Verdugo et al. 2019; Rossi et al. 2024), those of two other domesticated *Bos* species, namely the gaur (*B. gaurus*) and banteng (the continental *B. javanicus birmanensis* and the Indonesian *B. j. javanicus*), remain much more obscure. Domesticated banteng, known as Bali cattle (*B. j. domesticus*), is an economically important livestock in Southeast Asia, believed to be domesticated in present-day Indonesia from wild Javan banteng, which is also still extant as a wild species (Mohamad et al. 2009; Mohamad et al. 2012; Martojo 2012; Purwantara et al. 2012; Sudrajad et al. 2020). Here, we will use the terms ‘banteng’ for individuals descended from the Indonesian wild form, ‘Bali cattle’ for individuals descended from the domesticated form and *B. javanicus* as an umbrella term for both types. The *B. javanicus* domestication date remains contentious, and several studies report different times, e.g. around 5500 YBP (Rollinson 1994; Mohamad et al. 2009), as early as 7000 YBP (Felius 1995; Higham 2002; Lenstra et al. 2014), or 5000-10,000 YBP (Mohamad et al. 2012). Bali cattle have a restricted range in Southeast Asia, similar to the range of the banteng (Zhang et al. 2020), but here it is an important livestock representing more than 11 million individuals or approximately ∼27% of the total bovine livestock population in Indonesia (Mohamad et al. 2012; Purwantara et al. 2012). They are well adapted to small-scale village farming by being relatively small in stature (Kesuma 2019; Warmadewi 2020), resistant to several diseases, highly fertile, and possessing a remarkable ability to grow and produce milk on low-quality fodder (McCool 1992). Although Bali cattle are well adapted to the tropical climate and well-suited for the local husbandry conditions, it is not known to have been subject to intense selective breeding (Mohamad et al. 2012; Zhang et al. 2020). Originally found on the islands of Java and Bali, Bali cattle have recently been exported to other islands in Indonesia as well as surrounding countries such as Papua New Guinea, Malaysia, and Philippines (Lenstra et al. 2014). They were also introduced on the Cobourg peninsula in northern Australia (Bradshaw et al. 2006, 2007). Here, they established a large feral population descended from just 20 Bali cattle and subsequently released from a failed British outpost in 1849 (Bradshaw et al. 2006, 2007). This population now counts ca. 6000 individuals, making it the largest existing population of free-ranging *B. javanicus* in the world (Bradshaw et al. 2006, 2007).

Both taurine and zebu cattle were introduced into Indonesia during historical times (Leake 1980; Mohamad et al. 2009; Hartati et al. 2015; Sutarno and Setyawan 2015; Sudrajad et al. 2020), and were therefore brought into contact with *B. javanicus*. Crossbreeding between *B. javanicus* and cattle was widespread in Indonesia, but seems to always involve *B. javanicus* as the minor ancestry contributor (Wang et al. 2025 in review), and previous genetic studies found no evidence for introgression of cattle ancestry into Indonesian Bali cattle or banteng (Nijman et al. 2003; Mohamad et al. 2009, 2012; Wang et al. 2025 in review). Additionally, another study found no evidence of mtDNA or Y chromosomal allele sharing between either zebu or taurine with the Australian Bali individuals (Bradshaw et al. 2006, 2007). Nonetheless, genetic swamping of *B. javanicus* by the much more numerous cattle is considered a major threat to the species (The Zoological Record 1985; National Research Council and Research Coun National Research Council 2002; Mekong and Groenenberg 2024), and due to the lack of representative sampling this threat remains to be assessed.

The banteng has been assessed as Endangered on the IUCN Red List since 1996 and was uplisted to Critically Endangered in 2024 (Mekong and Groenenberg 2024). Over the past few decades, banteng have experienced range-wide population declines and restriction into small isolated subpopulations, most of which are still in decline, driven by human hunting for meat and trade in horns, combined with extensive loss and degradation of their habitat. For instance, there were six subpopulations of banteng on Java in 1990, but these were reduced to four by 2023 (Mekong and Groenenberg 2024). Due to very limited sample availability, the genetic characteristics of banteng and the history of domestication leading to Bali cattle is still largely unknown. Previous studies have largely focused on 1) coarse molecular markers such as mtDNA, Y chromosomal regions and microsatellites (Mohamad et al. 2009, 2012; Saijuntha et al. 2013; Qiptiyah 2019; Chaichanathong et al. 2021; Sun et al. 2022; Dagong et al. 2023), and 2) a few whole genome resequencing data mapped against the cattle reference genome instead of a banteng reference (Chen et al. 2018; Wu et al. 2018).

Here, we present a highly contiguous, chromosome level banteng genome assembly, along with whole-genome resequencing of Bali cattle from several locations, banteng bred in captivity, wild banteng and historical banteng samples. We first investigated the genetic structure and genetic diversity within *B. javanicus*, and inferred patterns of gene flow between cattle and *B. javanicus*. Next, we sought to establish the domestication history of *B. javanicus* and to investigate the demographic and selective forces acting during the domestication process. Our findings provide the first comprehensive investigation of one of the least publicized domestication events in human history. This will aid the effective management of genetic resources in the *B. javanicus* species complex, including in Bali cattle for commercial production, and can potentially inform conservation decisions regarding remaining wild *B. javanicus* populations.

## Results

### Assembling the first banteng genome

We sequenced, assembled and annotated the first reference genome for a banteng (*Bos javanicus*, RefSeq: GCF_032452875.1-ARS-OSU_banteng_1.0) using ∼60X long-read Oxford

Nanopore sequencing of an adult Javan banteng bull from a privately owned ranch in Texas, USA, assembled with Canu V2.1.1 (Koren et al. 2017). The final assembly consists of 1517 total scaffolds including 29 autosome-assigned scaffolds, sex chromosomes X and Y, and 1486 unplaced scaffolds spanning a total of 329 Mb (11% of total assembly). The total length of the assembly was 2.97 Gb, consistent with the K-mer estimated genome size, with a scaffold N50 of 87.5 Mb, and a scaffold N90 of 1.14 Mb (Figure 1A). It contains 12,777 out of 13,335 BUSCO orthologs (95.8%, Figure 1A; Figure S1; Table S1). A total of 35,317 genes, 21,582 protein coding regions, 60,389 mRNAs, 13,896 non-coding RNAs, and 60,628 CDSs were predicted by NCBI using the NCBI Eukaryotic Genome Annotation Pipeline, (see detailed description in https://www.ncbi.nlm.nih.gov/refseq/annotation_euk/Bos_javanicus/GCF_032452875.1-RS_2023_12/), consistent with a high quality assembly. We annotated nearly 54.63% of the genome as composed of repetitive elements, of which the majority are long interspersed nuclear elements (LINEs) (23.89%, Figure 1B; Table S2). Unplaced scaffolds were nearly entirely repetitive (98.43%) and mainly composed of Satellite repeats (90.54%; Table S2). We further identified that two samples from captivity in Texas, USA (LIB112408_Banteng_82B_Texas, LIB112405_Banteng_0805B_Texas) are first degree relatives of the individual used to generate this banteng assembly based on KING-robust kinship values > 0.200.

**Figure 1.**
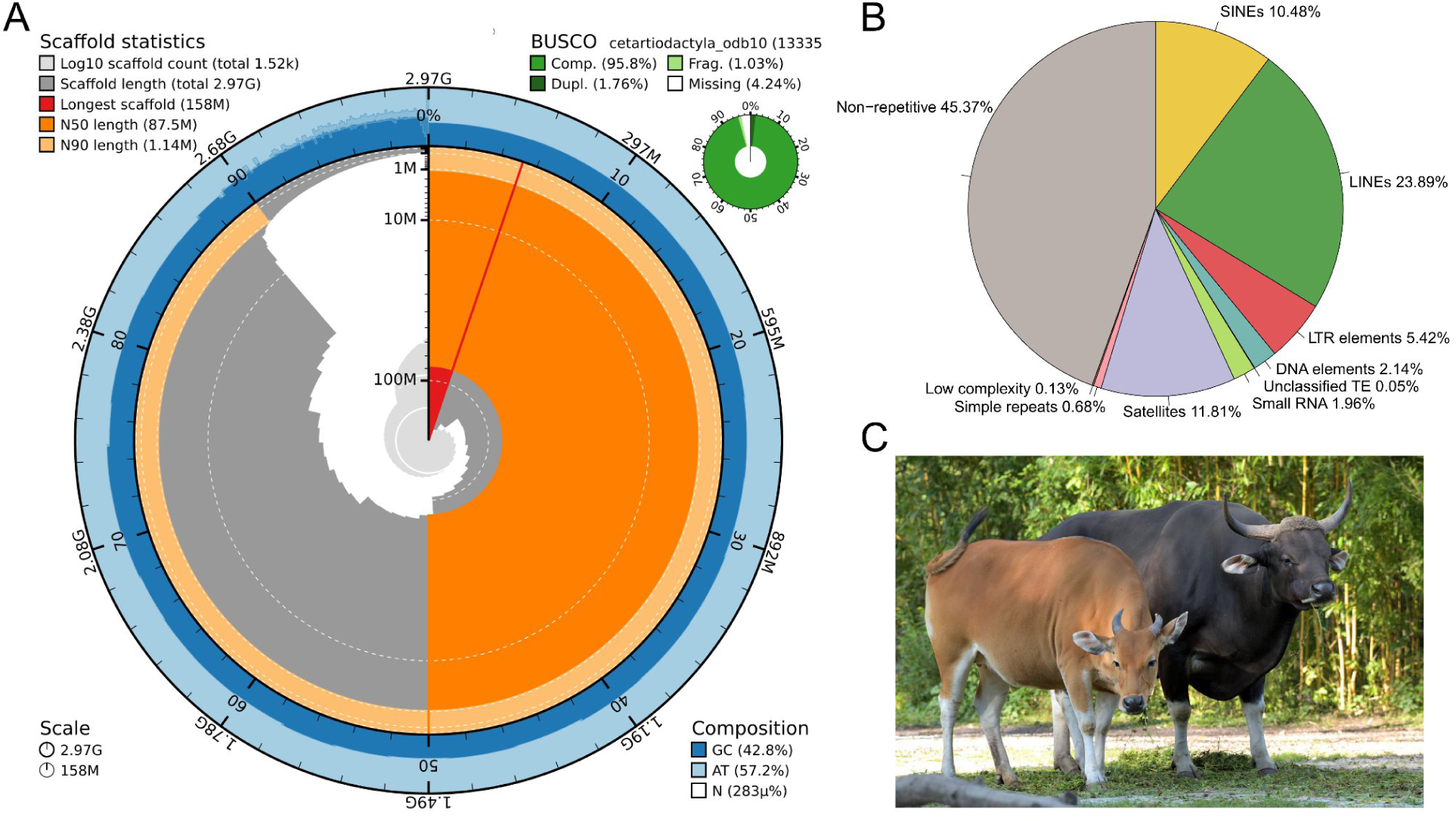
Summaries of assembly statistics. (**A**) Snail plot for banteng genome assembly. From outside to inside: dark blue and light blue rings show GC and AT percentages along the genome; the light orange segment represents N90 scaffold length; the orange segment represents N50 scaffold length; dark gray segments represent scaffold length distribution with plot radius scaled to the longest scaffold (red line); the central light grey shows log scaled scaffold count with white scale lines marking changes in order of magnitude. (**B**) Repetitive elements annotation. (**C**) Photograph of zoo-born banteng cow (left) and bull (right) from Hellabrunn Zoo, Munich, by Gemma Borrell (2020).

### Generating a population genomic dataset for *B. javanicus*, cattle and other bovines

We generated whole genome resequencing data for 19 Javan banteng (*B. j. javanicus*) samples, including 8 wild-born individuals currently kept in Indonesian zoos, 3 historical specimens collected in Java (between 1893 and 1905), and 8 specimens from a private ranch in Texas, USA. The samples were sequenced to a coverage ranging from 1.36X to 36.4X with a mean depth of ∼10.8X (Table S3). These genome sequences were supplemented with published whole genome sequencing data sets from five captive Javan bantengs, eight Bali cattle (*B. j. domesticus*) from Indonesia with no location of origin recorded (referred as Bali-Unk-Indonesia), and 46 Bali cattle previously generated by us (Wang et al. 2025 in review) from Bali (19), Kupang (15), and the feral population in northern Australia (12), respectively. We additionally downloaded 30 publicly available whole genome data sets representing other bovine species including 13 zebu (*B. t. indicus*), 10 taurine cattle (*B. t. taurus*), 4 gaur (*B. gaurus*), and 3 African buffalo (*Syncerus caffer*) as outgroups, leading to 108 individuals analysed in total (Table S3).

The sequence reads were mapped to three reference genomes, 1) the *de novo* banteng assembly presented here with a previously published mitochondrial genome added (GenBank: GCF_032452875.1-ARS-OSU_banteng_1.0; MT GenBank: JN632606.1), 2) the cattle reference genome *BosTau9* with a Y chromosome from a previous assembly added (GenBank: GCF_002263795.1-ARS-UCD1.2; Y chromosome GenBank: CM001061.2), and 3) a water buffalo (*Bubalus bubalis*) reference genome (GenBank: GCA_003121395.1-ASM312139v1). Initial sample quality filtering removed 7 individuals due to either extreme error rates and heterozygosity (1 sample), or relatedness (6 samples), resulting in 101 individuals for the final data set (Figure S2 and S3; Table S3 and S4). We identified 1,237,802,846 (41.6%) variant sites with reliable genotype calls in the banteng-mapped dataset, 1,152,481,983 (41.8%) in the BosTau9-mapped dataset, and 962,631,762 (36.2%) in the water buffalo-mapped dataset (Table S5) after rigorous filtering.

### Population structure and recent admixture

The population structure within *B. javanicus* populations including banteng and Bali cattle, and between *B. javanicus* and other bovine species, was first visualized by PCA using PCAngsd (Figure 2A; Figure S4 and S5). The first principal component (PC1) shows a clear separation of Australia Bali cattle from a cluster of Bali cattle from Bali, Kupang and Bali-Unk-Indonesia, whereas PC2 separates captive bantengs from the US from wild-born and historical bantengs (Figure 2A, lower panel). One captive banteng individual (B._javanicus_SD.Zoo_OR206 from San Diego Zoo, US, Heaton et al. 2016) is placed intermediately between the captive bantengs and the remaining individuals, indicating substructure or admixture in the lineages introduced to the captive population in US. A PCA of *B. javanicus* and other bovine species showed a slight affinity of Bali cattle from Kupang with the European taurine and South Asian zebu, confirming that Bali cattle from Kupang have been admixed recently with other cattle (Figure 2A, upper panel), as observed in Wang et al. (2025 in review). We also performed a PCA including the individual used to generate the banteng assembly, and found that it clusters with individuals from Texas, US (Figure S6).

**Figure 2.**
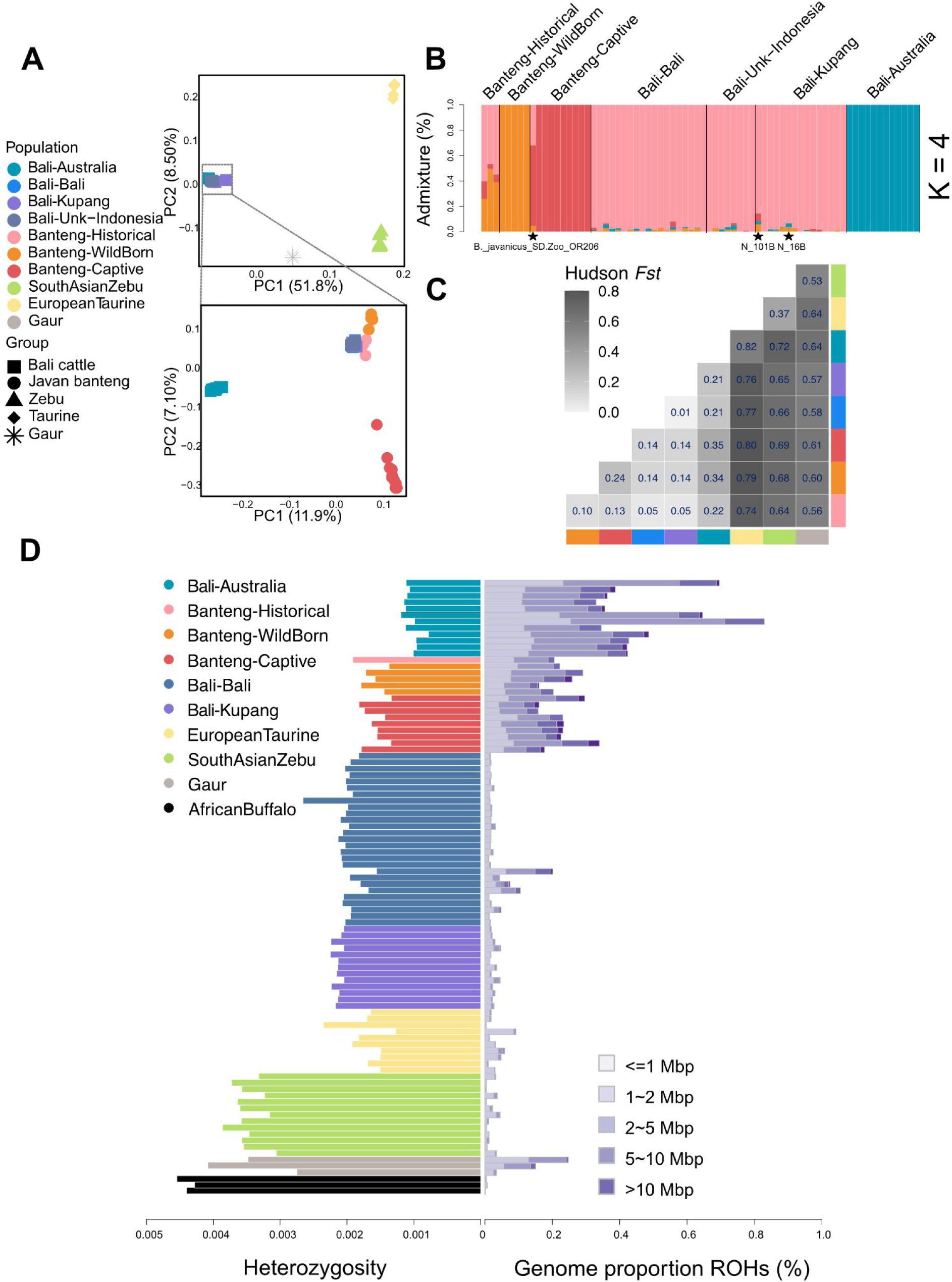
Population genomic analyses of banteng populations and other bovine species. **(A)** Principal component analysis showing genetic clustering among all individuals (upper panel) and within banteng populations (lower panels). **(B)** Admixture proportions of *B. javanicus* samples, estimated with NGSadmix assuming four ancestral clusters. Admixed individuals are indicated with their sample name and star symbol. **(C)** Genetic differentiation as described by pairwise Hudson’s *Fst* between pure defined populations. **(D)** Genome-wide heterozygosity and runs of homozygosity (ROH) proportions across each individual genome.

Admixture proportions based on NGSadmix further supported the PCA-predicted population structure (Figure 2B; Figure S7). The evalAdmix model with the highest number of converging ancestral source populations (K = 4) provides a good fit to the data without excessive correlation of residuals between populations (Figure S7), although some correlation of residuals within the inferred populations is still evident, suggesting some remaining substructure or distant relatedness among samples. Overall, individuals from wild-born banteng, captive banteng, Bali cattle from Australia, and the remaining Bali cattle from Bali, Kupang and from the unknown Indonesian locality, each formed an independent cluster, respectively, which is in line with the results of the PCA. Nine individuals, including the three historical bantengs, show potential signs of recent admixture, with their maximum admixture component lower than 0.95. However, only the two Bali cattle from Kupang (N_101B and N_16B), showed robust evidence of recent admixture as tested by the *apoh* method. The other seven samples show a very high inconsistency index in *apoh*, suggesting their admixture proportions are likely due to having ancestry from an unrepresented source, and should not be interpreted as recent admixture (Table S6).

Historical banteng, wild-born banteng, Australian Bali cattle, and Bali cattle from Bali and from the unknown Indonesian locality, were thus defined by the admixture analysis as four independent populations, hereafter referred to as ‘Banteng-Historical’, ‘Banteng-WildBorn’, ‘Bali-Australia’, and ‘Bali-Bali’, respectively. We removed the two recently admixed Bali cattle from Kupang from subsequent analyses and grouped the remaining Kupang individuals into their own population labelled as ‘Bali-Kupang’ due to their tentative evidence of introgression from cattle. We also discarded one individual (*B._javanicus*_SD.Zoo_OR206) from the captive banteng due to its unclear population assignment according to PCA and Admixture results, and grouped the remaining individuals into a population labeled as ‘Banteng-Captive’.

We quantified the genetic differentiation between the inferred banteng populations and found that *F*_ST_ is generally high among the three populations that separate on the first two PCA axes, Bali-Australia, Banteng-Captive and Banteng-WildBorn (*F*_ST_ = 0.24-0.35), whereas a very low *F*_ST_ was found between Bali-Bali and Bali-Kupang (*F*_ST_ = 0.01; Figure 2C), which is also consistent with the PCA result. Genetic differentiation between Banteng-WildBorn and the Bali cattle populations (Bali-Bali and Bali-Kupang) is 0.14. A NeighborNet tree based on the pairwise global *F*_ST_ distances conform with the PCA and admixture analyses, and suggests reticulation in the evolutionary relation between the populations (Figure S8).

### Genome-wide heterozygosity and recent inbreeding

We inferred the genome-wide heterozygosity for each sample except for two individuals from the Banteng-Historical population with low depth and high error rates (Figure 2D; Figure S2; Table S7). Overall, banteng and taurine individuals have similar median heterozygosities (0.0016-0.0019), which are clearly lower than zebu cattle (0.0036), gaur (0.0035), and African buffalo (0.0044). Domestication often reduces effective population size (*N*_E_) and genetic diversity (Ross-Ibarra et al. 2007; Bovine HapMap Consortium et al. 2009), however, we observed a slightly higher heterozygosity in Bali cattle (0.0011-0.0021) than in banteng, except for Australian Bali cattle (0.0011), consistent with its extreme recent founder event (Bradshaw et al. 2006, 2007).

Recent inbreeding and its effect on genetic diversity was investigated by identifying runs of homozygosity (ROH) in each individual (Figure 2D; Figure S9; Table S7). Bali-Australia has the highest proportion of ROH segments dominated by long ROH tracts (>5 Mb), suggesting inbreeding involving closely related individuals. All banteng individuals have much higher proportions of ROH than Bali-Bali, but lower than Bali-Australia. Similarly, Banteng-Captive has a larger proportion of ROH than Banteng-WildBorn. In contrast, Bali cattle (Bali-Bali and Bali-Kupang), taurine, zebu and African buffalo show low ROH proportions. We re-estimated heterozygosity in the genomic regions outside the ROHs to estimate heterozygosity without the influence of recent inbreeding and drift (Figure S10). Non-ROH heterozygosity was more uniform across banteng and Bali cattle populations, indicating that recent inbreeding and drift are the key drivers of the observed genetic variation across these populations. In contrast, genetic diversity in the other *Bos* species is more driven by other factors, such as ancient demographic history or the amount of admixture.

### Historical admixture between cattle and bantengs

The process of population divergence and inference of patterns of historical, as opposed to recent, admixture between populations was investigated using a TreeMix analysis on non-admixed individuals, allowing one migration edge (Figure 3A). The tree showed that all Bali cattle populations diverged from a single lineage within the *B. javanicus*. The inferred admixture event is from the South Asian zebu branch into Bali-Kupang.

**Figure 3.**
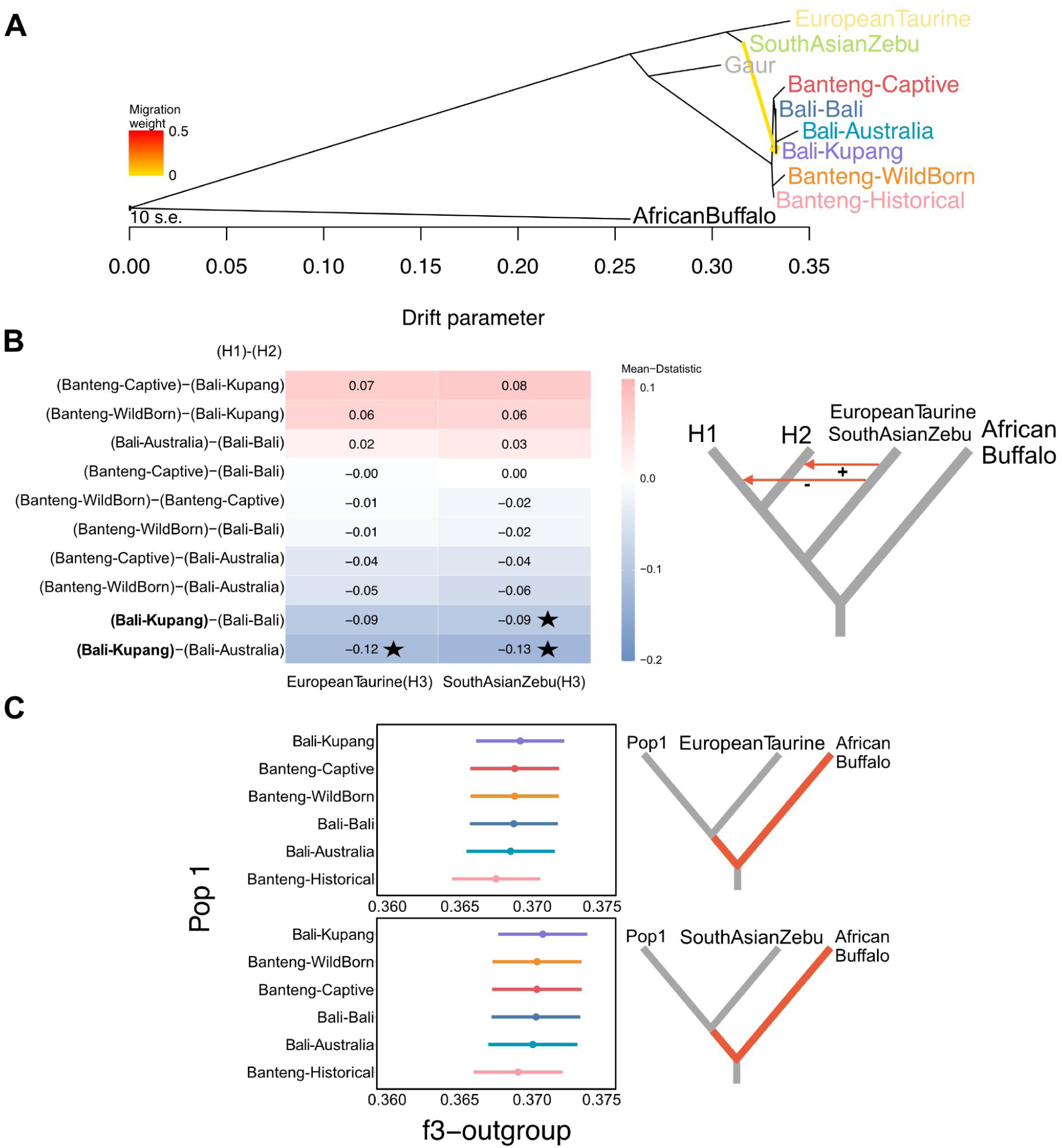
Historical admixture between Taurine/Zebu with Javan bantengs. **(A)** Population tree inferred with TreeMix assuming one migration event. Three African buffalo samples were used as an outgroup. The yellow arrow represents an inferred migration event, with colour indicating the migration weight (proportion of the admixed population estimated to derive from the source population). **(B)** D-statistics when using *B. javanicus* populations as H1 and H2, European taurine or South Asian zebu as P3 respectively, and African buffalo as outgroup H4, where the x-axis representing H3 and y-axis representing H1-H2. Star represents the significant median P value for each combination. D-statistics was estimated by single read sampling for all possible combinations of samples using ANGSD. The illustration to the right shows the interpretation of the sign of the D-statistic in the table on the left **(C)** Outgroup f3-statistics in the form f3 (X, taurine or zebu; African buffalo), where X denotes different *B. javanicus* populations.

D-statistic analyses confirmed that Bali-Kupang has a signal of excess allele sharing with South Asian zebu, although only significant relative to Bali-Australia and Bali-Bali (Figure 3B), and a significant signal of allele sharing with European taurine relative to Bali-Australia, indicating cattle introgression into Bali-Kupang after its divergence from other Bali cattle populations. Notably, this signal was found after removing two Bali-Kupang individuals with signals of recent admixture from cattle, suggesting cattle introgression into Bali-Kupang occurred both on a recent time scale where the Admixture method can detect it, and also further back in time.

We further investigated gene flow between cattle and *B. javanicus* populations using outgroup f3-statistics (Figure 3C). We observed that Bali-Kupang is genetically closer to cattle than the other *B. javanicus*, and that zebu cattle had a stronger genetic affinity with *B. javanicus* than taurine cattle, consistent with D-statistics analyses, although not significantly.

### Demographic history

Changes in historical effective population sizes were inferred using a Pairwise Sequentially Markovian Coalescent (PSMC) analysis on the non-admixed high-depth *B. javanicus* individuals (Banteng-Captive, Bali-Bali, Bali-Kupang and Bali-Australia), European taurine and South Asian zebu (five individuals for each population, Figure 4A). Both banteng and Bali cattle individuals exhibited similar demographic trajectories. The ancestral effective population size (*N*_E_) showed a peak at ∼1 million years ago (mya) followed by two distinct declines. The first decline occurred from ∼0.9 mya until ∼0.6 mya, coinciding with extensive multiple glaciation during the mid-Pleistocene in South Asia (Hope 2004). The second decline started from 40-50 thousand years ago (kya) in accordance with the beginning of the Last Glacial Maximum (LGM, De Deckker et al. 2003). European taurine cattle showed a similar demographic history, albeit with lower *N*_E_ after ∼200 kya, whereas the effective population size of South Asian indicine cattle increased after the divergence with taurine cattle.

**Figure 4.**
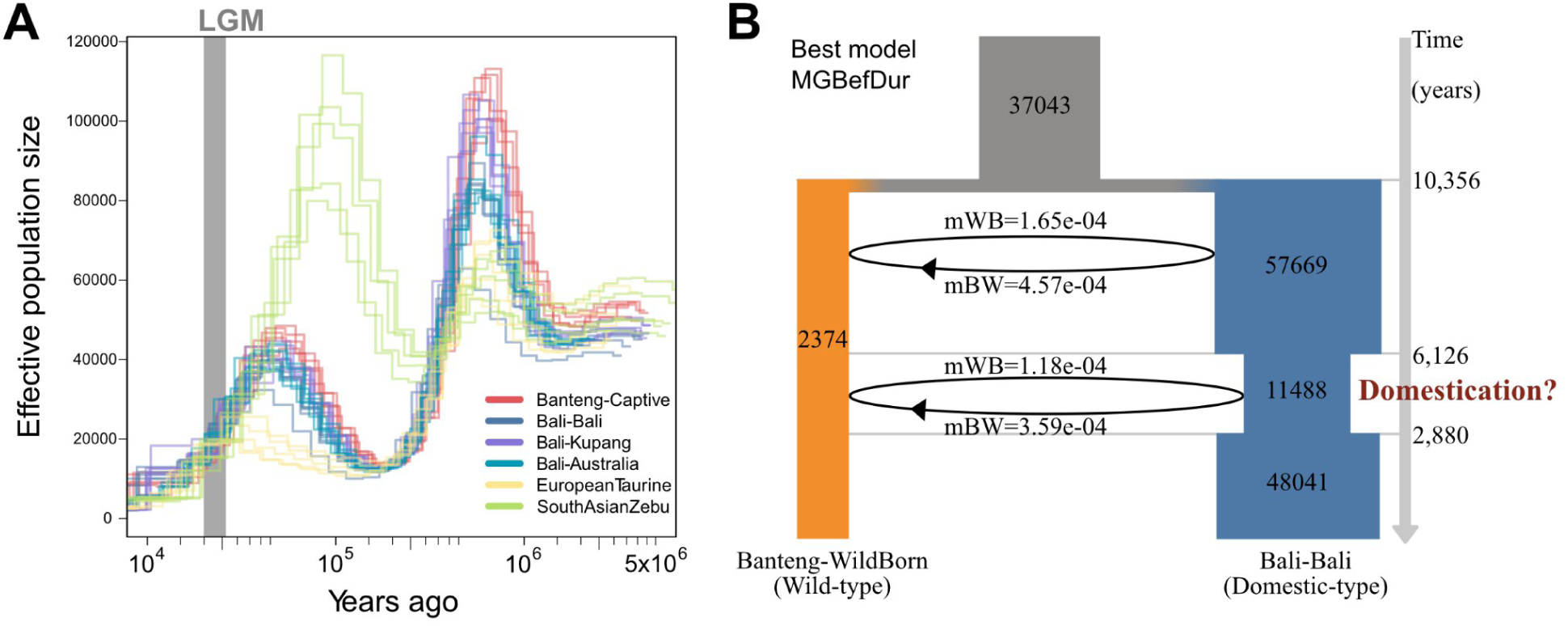
Demographic history of wild bantengs and Bali cattle. **(A)** Historical effective population sizes over time of different populations of *B. javanicus* and cattle using PSMC. **(B)** Schematic diagram depicting the best demographic model in Fastsimcoal2 using wild-born banteng and Bali cattle from Bali. Numbers within the bars are estimated effective population sizes. Inferred demographic parameters and 95% confidence intervals are shown in Table S9.

The joint demographic histories of banteng and Bali cattle were analyzed with two-dimensional site frequency spectra (2d-SFS) using Fastsimcoal2. The folded 2d-SFS was used to minimize potential bias in determining the ancestral allele (Figure S11), and Banteng-WildBorn and Bali-Bali was used as representative of a wild and a domesticated population, because the previous results suggested that these populations were the least influenced by recent bottlenecks, inbreeding and introgression. A set of 14 demographic models with different assumptions regarding i) the presence of a domestication bottleneck in Bali cattle, and ii) the presence of gene flow between banteng and Bali cattle, were initially evaluated (Figure S12). The “MGBefDur” model, a scenario with two periods of gene flow between banteng and Bali cattle (prior to and during the domestication bottleneck in Bali cattle), was the best-fitting model based on AIC and model-normalized relative likelihood, as recommended by Excoffier et al. (2013; Table S8). A comparison of estimated residuals between the simulated allele-frequency spectrum under the best model with the observed value from the actual data suggested that this model indeed captures the main features in the joint allele frequency distribution (Figure S13). Under this model, the split between the ancestor of extant banteng and Bali cattle was estimated to be 10,356 years ago, but with a large confidence interval (0-31,406; Figure 4B; Table S9). A moderate bottleneck in Bali cattle was inferred starting 4,999-7,253 years ago (CI) with a decrease in *N*_E_ from 57,669 to 11,488, lasting until 1,908-3,852 years ago (CI). We interpret this as the most plausible timing of the domestication of an ancestral banteng population, but we simultaneously note that any domestication bottleneck in Bali cattle—if present—was weak. Additionally, this analysis supported asymmetric gene flow pre-domestication and during domestication, with more gene flow from Bali cattle to banteng than in the other direction (Figure 4B; Table S9).

The divergence time between banteng and Bali cattle was also estimated using smc^++^, which does not allow gene flow between populations (Figure S14). Here, the population split was estimated to have occurred ∼7,800 years ago, which was regarded as a conservative lower bound on divergence time, as adding gene flow increases the divergence time estimates (Leaché et al. 2014). The two divergence time estimates based on either the SFS alone or the SFS plus linkage disequilibrium are therefore roughly consistent with each other.

### Selection during domestication

Genomic regions that may have been subject to selection during the domestication of *B. javanicus* were identified by scanning the genome for patterns of extreme divergence in allele frequency (elevated *F*_ST_), and extreme reduction in genetic diversity in the Bali cattle (π ln-ratio wild/domestic) in 50-kbp sliding windows between the two representative populations also used for the demographic analyses, Banteng-WildBorn and Bali-Bali. A total of 1148 outlier windows for *F*_ST_ and 483 outlier windows for the π ln-ratio based on a significance level of P-value <0.005 (Z test, with Fst >0.376 and π ln-ratio >1.33, Figure 5; Table S10) were identified after excluding any windows that contained <5,000 sites. We additionally calculated the XP-EHH statistics for each SNP between the same two populations and defined SNPs with a significant P-value < 0.005 (XP-EHH >2.81, Z test) as SNPs putatively under selection in Bali cattle. Overlapping outlier 50-kbp windows that were detected by at least two methods included: 65 windows shared by *F*_ST_ and π ln-ratio; 6 windows shared by *F*_ST_ and XP-EHH; 20 windows shared by π ln-ratio and XP-EHH, and 1 window shared across three statistics (Table S10). We merged outlier windows into a single region if the distance between windows was <1Mb, resulting in 49 outlier regions distributed across 19 chromosomes and ranging from 50 kbp to 550 kbp in size. The selection signal in these regions was further assessed by calculating Tajimas’ D and local linkage disequilibrium (LD) patterns in Bali-Bali (Figure S15).

**Figure 5.**
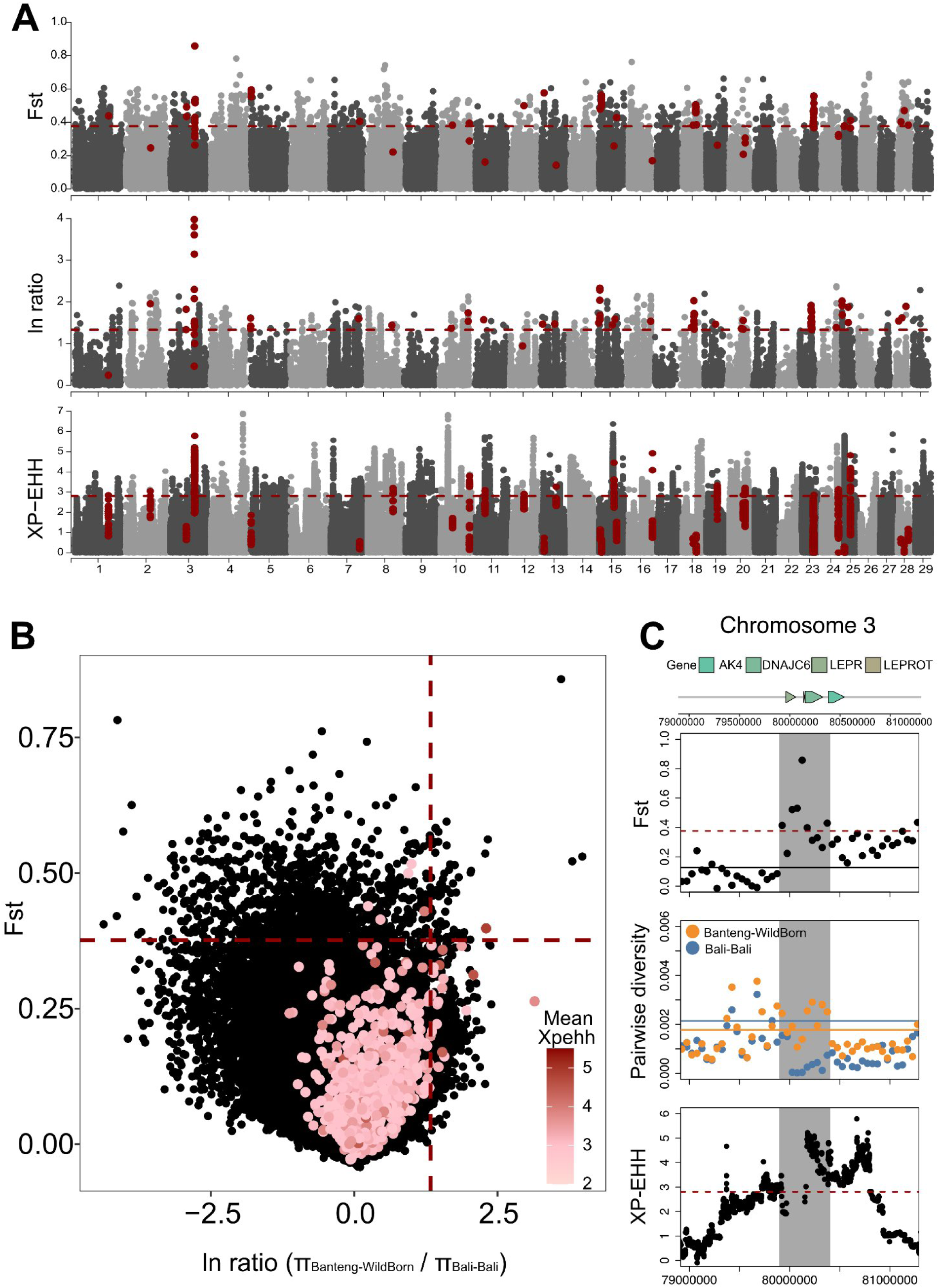
Genome-wide scans to detect genomic regions that have been subject to selection during domestication of banteng. **(A)** Manhattan plot for three selection scan statistics in 50 kbp windows across the genome: *F*_ST_, ln ratio (θπ,wild/θπ,domestic), and XP-EHH. Red dots mark windows that were outliers in two or more different scan statistics. **(B)** Distribution of the *F*_ST_, ln ratio (θπ,wild/θπ,domestic), and XP-EHH outlier values. The dashed vertical and horizontal lines indicate the significance thresholds (corresponding to Z test P<0.005, with *F*_ST_ > 0.376, π ln-ratio > 1.33 and XP-EHH > 2.81). **(C)** Example of outlier regions 79,900,000-8,0400,000 in chromosome 3 with selection sweep signals in Bali cattle. *F*_ST_, θπ and XP-EHH values and their annotated protein-coding genes are plotted. Horizontal solid lines represent the genomic mean of the corresponding statistics, dashed lines mark the outlier threshold. Grey shading indicates the inferred outlier region. Table S11 shows statistics of *F*_ST_, ln ratio, XPEHH, and annotated genes for all 49 regions that were contained in at least two selection scan outlier sets. Figure S15 shows plots of Tajima’s D and LD values for 49 regions.

These 49 outlier regions overlapped with 56 annotated genes associated with several physical characteristics, economically significant traits, reproduction and fertility traits, disease and development (Figure 5C; Table S10; Table S11). Only one gene present in at least two outlier sets is firmly linked to behavioral traits (*CDKAL1*), but two additional genes that were only present in the *F*_ST_ outliers (Z test, with *F*_ST_ > 0.376) are associated with aggression or behavioral traits (*BDNF, NTRK2*).

The only outlier region showing a selection signal across all three methods (79900000-80400000, *F*_ST_ = 0.451, π ln-ratio =2.45, mean XP-EHH=3.98) is located on chromosome 3 (Ref seq: NC_083870.1) and contains the genes *LEPR*, *LEPROT*, *AK4* and *DNAJC6* (Figure 5C). The leptin receptor (*LEPR* and *LEPROT*) genes are very well-known from human and animal studies and play important roles in fat deposition and energy homeostasis (Lu et al. 2007; Wylie 2011; Zhang et al. 2018; Almyah and Al-Badran 2019).

### Genetic load

The genetic load of the banteng and Bali cattle populations, as estimated by tolerant vs. intolerant substitutions using SIFT4G (Vaser et al. 2016), was compared to determine whether the domestication process led to an accumulation of harmful genetic variants. The exclusion of sites with transitions decreases the percentage of SIFT score by around two-fold, but does not change the relative difference of load between populations (Figure S16). We limit the load estimation to the samples with >4.5X, as we observed a negative correlation between the SIFT score and depth (Figure S16).

The banteng populations generally have lower total load than their domesticated relatives with significant statistical difference between Banteng-Captive and other populations (Figure 6A; ANOVA F=50.74, df=4, P<0.005; post-hoc Tukey all P<0.005). Banteng-Captive had the lowest total load of all populations (Figure 6A), whereas all the other populations showed similar value ranges. The trend persists whether we include or exclude ROH segments, as well as looking at only the homozygous genotypes (Figure 6B). In contrast, the single wild-born banteng with sufficiently high coverage to include in this analysis had higher total (Figure 6A) and homozygous (Figure 6B) genetic load than the other bantengs. However, low sample sizes for wild-born and historic banteng samples means that this conclusion is tentative at present.

**Figure 6.**
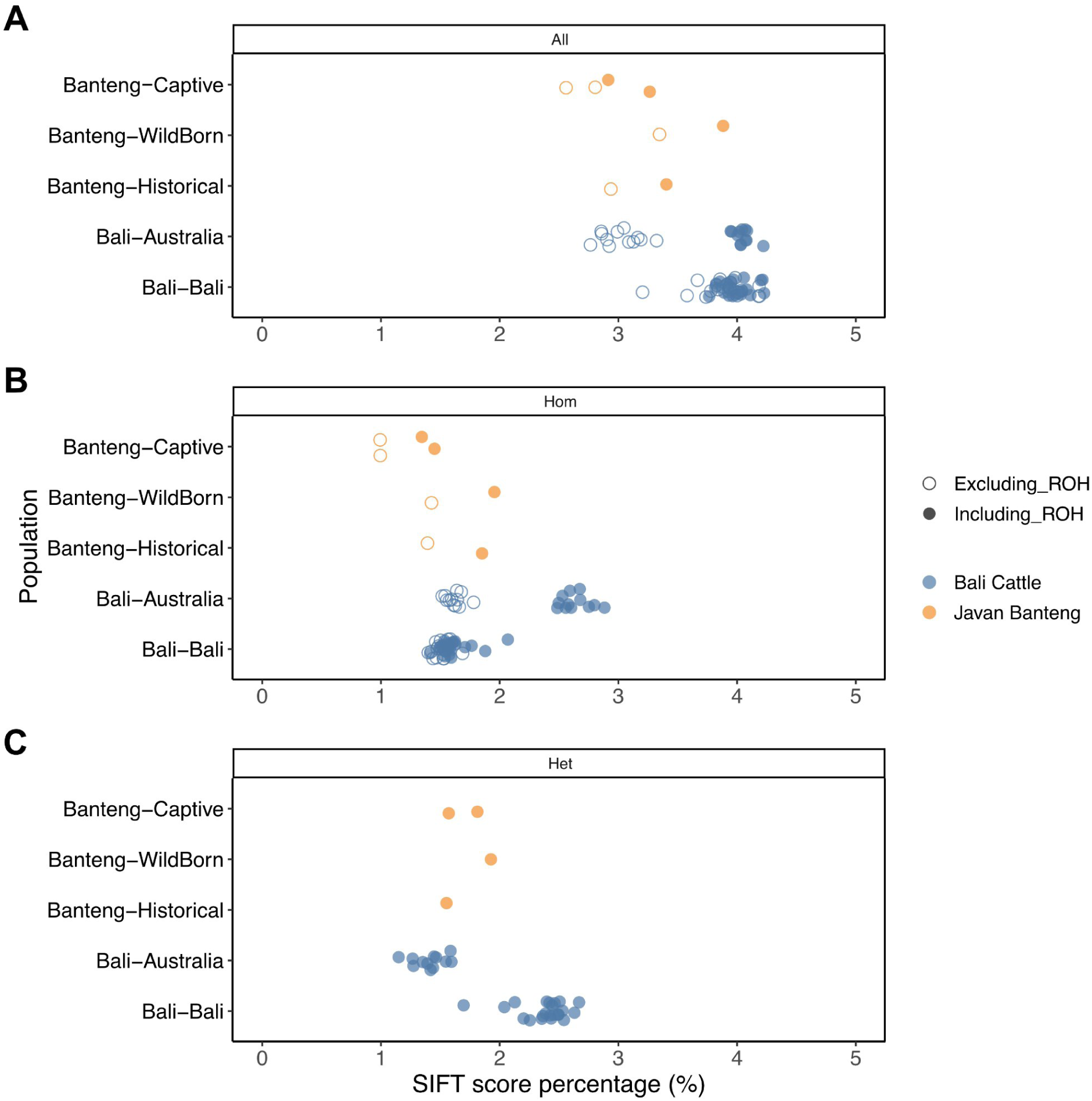
Genetic load measured by the percentage of SIFT score from maximum possible SIFT score in 66,548 deleterious sites (defined as SIFT < 0.05) separated between measurements on both homozygous and heterozygous genotypes. (**A**) only on homozygous genotypes (**B**) and only on heterozygous genotypes (**C**) accounting for the elevated load from including ROH segments (closed circle) compared to excluding ROH segments (open circle) in samples of comparable depth ranges.

Bali-Australia cattle were generally more inbred than other Bali cattle, although total genetic load was higher (4.05%) but not significantly different from Bali-Bali (3.98%, P=0.846). However, the distribution of SIFT score in homozygous and heterozygous state differed substantially between these two Bali cattle populations, with Bali-Australia having a much higher homozygous load score (mean score 2.64% vs 1.61%, P = <0.001), primarily caused by its high genomic proportion of ROH (Figure 6B). As many harmful mutations are believed to be recessive (Eyre-Walker and Keightley 2007), the homozygous SIFT score reflects the “realized load”, i.e. that experienced by the individuals at present (Bertorelle et al. 2022). On the other hand, Bali-Bali carries relatively more heterozygous load than Bali-Australia (2.38% vs 1.41%, P= <0.001, Figure 6C), which reflects the “potential load” because it can be expressed later in the population when recessive harmful mutations become homozygous (Bertorelle et al. 2022). Here, Bali-Australia had lower heterozygous SIFT score than all other populations (Figure 6C), possibly caused by genetic purging of harmful mutations caused by the extreme founder event and bottleneck experienced in this population.

## Discussion

### *B. javanicus* population structure is mainly driven by recent and man-made events

We generated the first high quality reference assembly of banteng, and provide new insights into the genetic basis of banteng domestication by performing a comprehensive whole-genome resequencing analysis of wild banteng and domestic banteng Bali cattle belonging to *B. javanicus*. Analysis of several whole-genome resequencing studies to characterize the genetic changes during domestication have compared domesticated populations with their wild progenitors, e.g. yak (Wang et al. 2014; Qiu et al. 2015), sheep (Zhang et al. 2022; Lv et al. 2022), goat (Dong et al. 2015; Zheng et al. 2020; Pogorevc et al. 2024), pig (Frantz et al. 2015), chicken (Wang et al. 2020, 2021) and reindeer (Wu et al. 2024). Here, we performed the first assessment of population structure in *B. javanicus* and found that most of the population structure is driven by recent extreme drift and inbreeding attributable to human intervention, primarily in Australian feral Bali cattle and captive bantengs (Figure 2A). This was supported by the highest *F*_ST_ being between the Bali-Australia and Banteng-Captive populations (*F*_ST_ = 0.35, Figure 2C), as well as extremely low heterozygosity and high ROH proportion in both populations (Figure 2D).

Disregarding the two highly drifted and inbred populations, we did not find evidence for strong population differentiation between banteng and Bali cattle. Genetic differentiation between Banteng-WildBorn and Bali-Bali was 0.14 (Figure 1C), which is smaller than *F*_ST_ between divergent taurine cattle breeds (*F*_ST_ = 0.215, Porto-Neto et al. 2014), and between dogs and wolves (*F*_ST_= 0.223, Axelsson et al. 2013). However, it is higher than *F*_ST_ between wild and domestic yaks, which have experienced continuous gene flow, including extensive post-domestication gene flow (*F*_ST_ =0.058, Qiu et al. 2015), and similar to differentiation between wild and domestic reindeer (*F*_ST_ = 0.062-0.145), a domestication that was characterized by weak artificial selection pressure and an absence of a strong domestication bottleneck (Wu et al. 2024). We also found that the demographic history of banteng and Bali cattle is indistinguishable until the recent past, and that heterozygosity across all *B. javanicus* was more uniform when removing ROH regions from the heterozygosity estimation (Figure S10). These observations suggest that all extant *B. javanicus* share the vast majority of their evolutionary history, and that there is little ancestral population structure in this species. We do note, however, that the limited availability of wild Javan banteng samples preclude firm conclusions regarding the species-wide population structure, and future work including samples from isolated populations across Java are important to resolve this question.

### Banteng domestication history: a weak bottleneck, substantial gene flow and low introgression

We found that, beyond the particular cases of extreme drift in the feral Australian population and the Banteng-Captive population, banteng in general have low to moderate genetic diversity, comparable to taurine cattle but lower than zebu, gaur and African buffalo. Demographic history inference suggested two Pleistocene bottlenecks, of which the most recent one coincides roughly with the onset of the Last Glacial Maximum (Figure 4A, De Deckker et al. 2003). Other bovine species such as yak (Qiu et al. 2015) and taurine cattle (Chen et al. 2018) living in South Asia also suffered decreases in *N*_E_ during the same periods, suggesting a possible regional disturbance associated with the Pleistocene glaciations (Hope 2004). Consistent with the lack of ancestral *B. javanicus* population structure, we found that the divergence between the ancestral banteng and Bali cattle populations was relatively recent (10,356 YBP) and that gene flow persisted between the two for thousands of years after this divergence. Notably, the long-term effective population size of the lineage eventually leading to Bali cattle was substantially larger than that of the wild population after this split, suggesting that domestication occurred from a larger banteng subpopulation than the one directly ancestral to extant wild banteng. In addition, we found no support for a strong bottleneck in the domesticated lineage, which was only reduced to an effective size of ∼11,000 individuals at its lowest point between ∼6000-2900 YBP, which therefore represents our best estimate of a domestication time frame, although we emphasize that the lack of a clearly defined bottleneck makes it hard to ascertain when the lineage leading to Bali cattle was actually or fully domesticated in its present form. Interestingly, these factors together have led to a higher current genetic diversity in Bali cattle than in banteng, a pattern shared by other domesticated animals such as yaks (Qiu et al. 2015) and pigs (Ojeda et al. 2006; Iacolina et al. 2016). In these species, sustained gene flow with their wild relatives appears to play an important role in explaining their relatively high genetic diversity. We also found post-divergence gene flow between banteng and Bali cattle, albeit with more gene flow going into banteng than the reverse, and a cessation of gene flow in recent times (Figure 4B). In *B. javanicus*, wild-domestic gene flow has not led to high genetic diversity, possibly because the two lineages were never highly differentiated from each other. Overall, our results point to the domestication of banteng being a gradual process that may have occurred over millennia and involved continuous sourcing of wild bantengs into the domestic populations, and that the evolutionary divergence between the two is subtle. On this evidence the domestication of *B. javanicus* appears more similar to the gradual and “soft” domestication of either yak or reindeer than to other well-known animal domestications. This is consistent with the observation of limited morphological differentiation between the two forms (Felius 1995; Higham 2002; Lenstra et al.2014), in contrast with most other co-existing wild/domesticated animal forms (Diamond 2002; Carneiro et al. 2014; Ahmad et al. 2020).

In contrast, introgression from cattle appears to be very limited in *B. javanicus*, with the exception of the Bali cattle population from Kupang, in which we found indications of both recent (Figure 2) and older (Figure 3) introgression. This cattle introgression is also reflected in higher heterozygosity compared with other Bali cattle populations (Figure 2D), but our analyses do not support major cattle introgression, as the D-statistics are low and only borderline significant. This is in contrast with Malaysian Bali cattle that showed evidence of larger-scale zebu introgression, with 65% of mtDNA haplotypes in the population being of cattle origin (Nijman et al. 2003). Hybridization or introgression can lead to extinction of populations or species by genetic swamping (Rhymer and Simberloff 1996; Allendorf et al. 2001; Todesco et al. 2016), a process in which the parental lineages are gradually replaced by hybrids (Rhymer and Simberloff 1996; Allendorf et al. 2001; Todesco et al. 2016). Taken together, these and previous findings demonstrate that despite the ability of banteng and cattle to hybridize and the presence of highly admixed cattle populations, introgression is not at present a major concern for the genetic coherence of *B. javanicus* as a whole, alleviating both conservation concerns (The Zoological Record 1985; National Research Council and Research Coun National Research Council 2002; Mekong and Groenenberg 2024) and economical concerns of genetic swamping of Bali cattle (Martojo 2012).

### Selection signatures despite the “soft” domestication history

The lack of a strong domestication bottleneck and the presence of gene flow post-domestication reduces the expectation of finding genes under strong selection during *B. javanicus* domestication. Despite the weak signal of a domestication bottleneck in Bali cattle, we identified 56 candidate genes under positive selection during domestication. The leptin receptor gene *LEPR*, a gene located on the only outlier region across three selection scan methods, is well-known for association with body weight, lipid composition, feed intake and metabolism traits in a wide range of species, and also with reproduction and fertility, possibly as an indirect consequence of the metabolism association (Wylie 2011). It has been found to be associated with obesity and type 2 diabetes in humans (Lu et al. 2007; Zhang et al. 2018; Almyah and Al-Badran 2019). In animals, the effects of *LEPR* on economically important traits have been widely investigated, including growth traits in Chinese indigenous cattles (Guo et al. 2008), fat and protein contents in Jersey cattle (Komisarek and Dorynek 2006), reproduction traits such as calving interval in Slovak Spotted and Pinzgau cows (Trakovická et al.2013), the age at first insemination in Polish cattle (Komisarek 2010), the ability to recycle after calving (ICF) in Chinese Holsteins (Zhang et al. 2023), and seasonality reproduction in Rasa aragonesa sheep (*Ovis aries*, Lakhssassi et al. 2020), and both growth and carcass traits in Savimalt and French Giant meat-type quails (Wang et al.2023). Additionally, among the genes that were outliers in at least two selection scans were genes associated with potentially production-relevant traits such as carcass traits and meat quality, body conformation and growth, feed efficiency, milk production, fertility and disease (Table S11). This list also includes genes associated with typical domestication traits such as coat color and heat adaptation (Table S11). However, despite our finding that these genes are potentially under selection in *B. javanicus*, the biological significance of variants in these genomic regions is unknown.

Despite reports that Bali cattle are more docile than banteng (Martojo 2012), which is consistent with an expectation that behavior and tameness are among the most crucial domestication traits (e.g. Jensen 2014; Agnvall et al. 2018; Katajamaa and Jensen 2021), we only found one gene known to be related to behavioral traits to be supported across two selection scan methods: the neurotransmitter concentration regulator *CDKAL1* (Chen et al. 2020). Several additional behavior-related genes were present in the *F*_ST_ outliers (*F*_ST_ > 0.376), but were not corroborated by the other scan statistics. One such gene, the brain-derived neurotrophic factor *BDNF* is highly expressed in the brain and is a candidate gene for psychiatric disorders such as Bipolar Disorder (Grande et al. 2010), and schizophrenia (Zhang et al. 2016) in humans, and associated with memory and aggression in rats (Moskaliuk et al. 2023). The high-affinity receptor of *BDNF* is encoded by another gene present among the *F*_ST_ outliers, neurotrophic tyrosine kinase receptor type 2 *NTRK2*.

### Implications for managing the genetic resources in banteng

Conserving the genetic diversity of *B. javanicus* is of imminent importance considering their Critically Endangered status. While banteng populations generally have reduced heterozygosity compared to their domesticated relatives (Figure 2D), they have a lower inferred genetic load (Figure 6). Also of note, we found that genetic diversity and load in banteng have not changed substantially since 1893-1905, the time of sampling of the historical samples included here. The higher total load in Bali cattle is consistent with a typical domestication syndrome where human husbandry results in relaxed selection pressure, so that deleterious mutations accumulate more readily (Larson and Fuller 2014; Wang et al. 2011). The lower total load in banteng, however, does not necessarily mean that the population is less susceptible to future changes. Low genome-wide heterozygosity is associated with lower evolutionary and adaptive potential (Kardos et al. 2021; DeWoody et al. 2021), and it may also be significant that the only wild-born banteng in which we were able to estimate genetic load actually had load on a similar level as the Bali cattle (Figure 6). We support previous conclusions that the feral Bali cattle population in Australia, despite the fact that it is presently the largest wild population of *B. javanicus*, should not be a first choice reservoir of the *B. javanicus* gene pool as it is highly inbred (Wang et al. 2025 in review and this study), and we furthermore corroborate this by showing that it has high homozygous load. We also highlight that other *B. javanicus* populations, notably Bali cattle from Bali, carry substantial potential, or masked, load that can become realized if the populations become more inbred or experience a bottleneck, as exemplified by the Australian Bali cattle. Direct observations in the Australian feral populations, compared with other populations, would be useful to reveal whether the increased inbreeding and higher homozygous load translates into observable fitness loss. Finally, our results show that Javan banteng have limited genetic structure, even when considering wild and domesticated populations as a whole. Hence, the relative genetic uniformity between domestic and wild Javan banteng could be important for future decisions regarding the management of the subspecies’ genetic resources.

Our study represents the first comprehensive study of *B. javanicus* on a genomic scale, as well as a high-quality banteng reference genome that is likely to find wide use for studying *Bos* evolution. Due to the limited number of wild-born banteng samples included here, our conclusions are by necessity restricted. We call for the generation of additional sequencing data representing not only banteng, but from severely underrepresented wild *Bos* in general.

## Methods

### Genome assembly and annotation

#### Assembly and scaffolding

High molecular weight DNA from blood was prepared for long read sequencing on the PromethION instrument with Ligation Sequencing Kit LSK-110 (Oxford Nanopore Technologies, Oxford, U.K.). Fastq files were generated from fast5 raw data using the Guppy basecaller v4.4.2 (https://nanoporetech.com/software/other/guppy/history?version=4-4-2) with the bonito research model v0.3.2 (res_dna_r941_min_crf_v032.cfg). A total of 4 flow cells were collected yielding 12,529,067 reads totaling 278,721,464,370 bp >Q7 with a 41,833 bp N50. The same DNA sample was used to generate a short-read library using the TruSeq PCR-Free DNA kit (Illumina Inc., San Diego CA) and sequenced on a NovaSeq with 150 bp paired-end reads. Long-read sequence data was assembled using Canu v2.1.1 with parameters “ -nanopore ‘corMhapOptions=--threshold 0.8 --ordered-sketch-size 1000 --num-hashes 512 --ordered-kmer-size 14’ ‘correctedErrorRate=0.105’ ‘corMhapSensitivity=normal’ ‘corOutCoverage=100’ ‘MhapBlockSize=1500’ ‘genomesize=2.7g’”. Assembled contigs were polished with long reads using two rounds of PEPPER-Margin-DeepVariant v0.4 variant calling (https://www.nature.com/articles/s41592-021-01299-w) with bcftools (https://doi.org/10.1093/bioinformatics/btp352) consensus generation. Purge_dups v1.2.5 (https://doi.org/10.1093/bioinformatics/btaa025) was used to remove partially duplicated and low-coverage contigs, likely representing errors, to generate a final set of contigs.This was followed by two rounds of short read polishing using Freebayes (version v1.3.1-1; https://arxiv.org/abs/1207.3907) with bcftools consensus generation. Variants called for polishing with both methods were screened with Merfin (https://github.com/arangrhie/merfin, version downloaded October 20, 2020) which predicts the k-mer consequences of variant calls and validates supported variants. The final contigs were ordered and oriented through alignment with the cattle genome *BosTau9* (GenBank: GCF_002263795.1-ARS-UCD1.2) using minimap2 with the parameter “-x asm10”.

#### Gene annotation, repeats annotation and assembly evaluation

Annotation of banteng reference genome (*Bos javanicus*, RefSeq: GCF_032452875.1-ARS-OSU_banteng_1.0) were automatically performed by NCBI using the NCBI Eukaryotic Genome Annotation Pipeline, an automated pipeline that annotates genes, transcripts and proteins on draft and finished genome assemblies. All of annotation products are available in the sequence databases in NCBI (https://ftp.ncbi.nlm.nih.gov/genomes/all/annotation_releases/9906/GCF_032452875.1-RS_2023_12/). We used RepeatMasker V4.1.1 (http://www.repeatmasker.org/) to identify repeat regions in the reference genomes, utilizing ‘rmblast’ as the search engine and ‘mammal’ as the query species with default settings. To assess the quality of genome assembly, we calculated assembly statistics by blobtools V1.1.1 (Laetsch and Blaxter 2017), evaluated completeness using lineage dataset ‘cetartiodactyla_odb10’ by BUSCO V5.4.4 (Simão et al. 2015), and compared statistics with the other two references used in this study *BosTau9* with Y chromosome (GenBank: GCF_002263795.1-ARS-UCD1.2; Y chromosome GenBank: CM001061.2), and *Water buffalo* (GenBank: GCA_003121395.1-ASM312139v1). We also investigated the closely relatedness and population structure between banteng assembly and the other individuals in this study.

### Sampling and lab protocol

We collected 19 samples of banteng (*B. j. javanicus*) in total, out of which eight are wild born Javan banteng, three historical Javan banteng specimens dated to 1893-1905 years, and 8 captive Javan banteng from a captivity in Texas, USA. Collected blood samples were kept in a DMSO buffer in the field, stored at -20 °C as soon as possible, and were further transferred to a -80 °C freezer for long-term storage. Following the manufacturer’s protocol instructions of the QIAGEN Blood and cell culture Kit, we performed DNA extraction. Due to the humid climate in the Indonesian lab, we added three extra treatment steps before following the default protocol: (1) adding 500 µl of ice cold water to the blood samples, (2) centrifuging the diluted blood samples for 20 min 4000 rpm in 4°C, and (3) discarding the supernatant without disturbing the pellet. DNA concentrations were further measured with a Qubit 2.0 Fluorometer and a Nanodrop, and the quality of genomic DNA was checked by gel electrophoresis. After DNA extraction, we fragmented 1 mg genomic DNA by Covaris (350 bp on average), and performed purification using AxyPrep Mag PCR clean up kit. The fragments were then end-repaired by End Repair Mix and were purified. We then combined the repaired DNA with A-Tailing Mix, and ligated the Illumina adaptors to the DNA adenylate 3’ ends, followed by product purification. Size selection was performed targeting insert sizes of 350 base pairs. We finally performed several rounds of PCR amplification with PCR Primer Cocktail and PCR Master Mix to enrich the adaptor-ligated DNA fragments. After purification, the size and quality of libraries were assessed by the Agilent Technologies 2100 Bioanalyzer and ABI StepOnePlus Realtime PCR System.

We additionally downloaded published whole genome sequencing data sets from five captive Javan bantengs (*B. j. javanicus*), eight Bali cattle (*B. j. domesticus*) from Indonesia with no specific location of origin recorded, and 46 Bali cattle (*B. j. domesticus*) previously generated by us (Wang et al. 2025 in review) from Bali (19), Kupang (15), and northern Australia (12), respectively. Bali cattle from Australia are part of a feral population in Garig Gunak Barlu National Park in northern Australia, which descended from 20 individuals that were released from a failed British outpost in 1849 (Bradshaw et al. 2006, 2007). Moreover, 30 publicly available whole genomes representing other bovine species including 13 zebu (*B. t. indicus*), 10 taurine cattle (*B. t. taurus*), 4 gaur (*B. gaurus*), and 3 African buffalo (*Syncerus caffer*) as outgroups were also downloaded, leading to 108 individuals in total for mapping and subsequently samples/sites filtering (Table S3).

### Sequencing and mapping

Small pieces of damaged loose bone were collected from three historical Javan banteng skulls specimens, from the collections of the Natural History Museum of Denmark. Collection ID’s: CN 634 (collected 03-12-1868), CN 904 and CN 906 (both collected 08.07.1905). The bone samples were processed under strict clean laboratory conditions at the Globe Institute, University of Copenhagen. The bone pieces were manually cut, then placed into 2 ml microcentrifuge tubes and washed with a 7% bleach solution, ethanol and ddH2O, following Boessenkool et al. (2017). After washing, samples were pre-digested for 15 minutes at 37 °C with rotation, in 1 ml of freshly made digestion buffer containing 1M Tris-HCL, 5M Sodium chloride, 1M Calcium chloride, 0.5M EDTA, 10% SDS, 1nM DTT and 10 µg/µl of proteinase K, following Gilbert et al. (2007) DNA extraction protocol. The pre-digestion lysate was discarded, after which a second digestion with 1 ml of the same buffer and settings was performed overnight. After incubation, tubes were centrifuged on a bench-top centrifuge at 6,000 x g for 3 minutes. The lysate was then additionally treated with phenol-chloroform clean up following Carøe et al. (2018). The supernatant was then purified using MinElute column with modified PB buffer and eluted using 2 washes in pre-warmed 18 μL buffer EB (Qiagen) - with 3 min of incubation time at 37°C (Dabney et al. 2013). The DNA concentration of each extract was verified on a Qubit (ng/μl). 16.25μl of purified DNA was treated with 5 μl of (1U/1μl) USER™ enzyme (New England BioLabs®, Inc.) and incubated for 3 hours at 37°C, followed by Blunt-End dsDNA library preparation (Meyer and Kircher 2010), following modifications in Gamba et al. (2014). AccuPrime Pfx DNA Polymerase (Invitrogen) was used for the PCR amplification with 3μl of DNA library and PCR cycles varying from 8-12 cycles. PCR products were purified using minElute columns (Qiagen) and were quantified using a Tapestation 2200 (Agilent technologies). Indexed PCR libraries were sequenced on to 1.36X-5.61X coverage on an Illumina NovaSeq6000 platform (Illumina Inc.). Modern samples were sequenced using Illumina paired-end 2×150 bp reads. This includes 8 wild born Javan banteng samples sequenced to medium depth of 4.32X-5.17X coverage on the Illumina NovaSeq6000 platform, 8 samples from captive Javan banteng sequenced to high depth of 12.6X-36.4X coverage on the Illumina HiSeq2500 platform (Illumina Inc.).

We evaluated the quality of the raw reads using FastQC (bioinformatics.babraham.ac.uk/projects/fastqc) and MultiQC (Ewels et al. 2016) before mapping. We used the PALEOMIX BAM pipeline to perform mapping (Schubert et al. 2014). We first trimmed Illumina universal adapters using AdapterRemoval v2.3.2 (Schubert et al. 2016). To improve the fidelity of the overlapping region by selecting the highest quality base when mismatches are observed, we then merged read pairs with overlapping sequences of at least 11 bp. Mismatching positions in the alignment, where both read bases had the same quality were set to ‘N’ via the ‘--collapse-conservatively’ option. No trimming of Ns or low-quality bases was performed and we only discarded the empty reads resulting from primer-dimers. All trimmed reads were then subsequently mapped using BWA-mem v0.7.17-r118870 (Li 2013) to three chromosome-level reference genomes: 1) *de novo banteng* assembly presented here with MT genome (GenBank: GCF_032452875.1-ARS-OSU_banteng_1.0; MT GenBank: JN632606.1), an adult bull banteng from privately owned ranch in Texas, USA, (2) *BosTau9* with Y chromosome (GenBank: GCF_002263795.1-ARS-UCD1.2; Y chromosome GenBank: CM001061.2), a female taurine from Hereford breed, and (3) *Waterbuffalo* (GenBank: GCA_003121395.1-ASM312139v1), a female water buffalo from the Mediterranean breed. We flagged PCR duplicates using samtools v1.11 ‘markdup’ for paired reads and PALEOMIX ‘rmdup_collapsed’ for merged reads. The resulting BAM alignments from collapsed and paired reads were merged for each individual, and filtered based on standard BAM flags to exclude unmapped reads, reads with unmapped mate reads, secondary alignments, reads that failed QC, PCR duplicates, and supplementary alignments. Further, reads in alignments with inferred insert sizes smaller than 50 bp or greater than 1000 bp were excluded, reads where fewer than 50 bp or less than 50% of the read were aligned, and read pairs in which mates mapped to different contigs or not in the expected orientation were also discarded. Finally, we generated statistics of the filtered BAM files by samtools ‘stats’ and ‘idxstats’ (Li et al. 2009).

### Sample filtering

#### Assessing post-mortem damage in historical samples

For historical samples, we used mapDamageV2.0 (Jónsson et al. 2013) to assess the post-mortem in samples, as fragmentation and base misincorporation are the main characteristics of damaged aDNA molecules. We didn’t observe any surplus of C-to-T misincorporations at the 5’ ends of sequences and complementary G-to-A misincorporations at the 3’ termini, which are caused by increased cytosine deamination in single-stranded 5’-overhanging ends (Figure S2A). We thus kept all of three historical samples for downstream analyses.

#### Error rates

For modern samples, we identified and removed samples with excessive sequencing error rates using the ‘perfect individual’ method (Orlando et al. 2013) implemented in ANGSD. The rationale behind this approach is that each ingroup sample should have the same expected number of derived alleles as the ‘perfect individual’ when using an outgroup, and that a surplus of observed derived alleles in each sample is thus due to excess errors relative to the ‘perfect individual’. We first built the consensus sequence for the ‘perfect individual’ (a50) using the ‘-doCounts 1 -doFasta 2’ option in ANGSD, and then we estimated error rates using all bases (-doAncError 1), while both steps setting the quality filters of a minimum base quality of 30 (-minQ 30) and a minimum mapping quality of 25 (-minMapQ 25). Based on this, one sample of CaptiveJava (B._javanicus_W_Zoo_DB) from UK Zoo showing extremely high error rates (> 0.001) was removed for downstream analyses (Figure S2B-C; Table S3).

#### Heterozygosity

For modern samples, we next excluded samples with extraordinarily high levels of heterozygosity as these samples very likely suffer from DNA contamination or considerable sequencing errors. To calculate individual heterozygosity, we estimated one-dimensional site frequency spectrum (1d-SFS) with genotype likelihoods using the GATK model in ANGSD. The analysis revealed one individual with excessively high heterozygosity (∼0.00570), which is the same sample identified by extremely high error rates, thus was excluded for downstream analyses (Table S3).

#### Relatedness filtering

We identified closely related and potentially duplicated samples based on KING-robust kinship coefficient described in Waples et al. (2019), which can be used to detect close familial relatives without estimating population allele frequencies. To calculate KING-robust statistics, we estimated two-dimensional site frequency spectra (2d-SFS) for individual pairs with genotype likelihoods using the GATK model (-GL 2) in ANGSD. We identified six pairs of first degree relatives based on KING-robust kinship values > 0.200. For each of these pairs, we processed to exclude all but one sample with lower coverage from identified related pairs, leading to five samples being discarded (Table S3; Table S4).

Before downstream analyses, we performed a second and more sensitive analysis of relatedness within each sampling location by using NgsRelate, which can be used to infer relatedness for pairs of individuals from low coverage Next Generation Sequencing (NGS) data by using genotype likelihoods instead of called genotypes (Korneliussen and Moltke 2015); (Hanghøj et al. 2019). We identified one additional pair of relatives within the sampling localities of gaur and removed the lower depth sample ‘gaurus_O_Zoo_5681’ (Figure S3). Finally, 101 out of 108 samples were retained after sample filtering.

### Sites filtering

#### Reference genome filtering

We implemented multiple filtering for three reference genomes based on different criteria. We used RepeatMasker V4.1.1 (http://www.repeatmasker.org/) to identify repeat regions in the reference genomes, utilizing ‘rmblast’ as the search engine and ‘mammal’ as the query species with default settings. GenMap V1.2.0 (Pockrandt et al. 2020) was used to calculate the mappability score of each site, conservatively using 150 bp k-mers with up to two mismatches allowed (-K 150 -E 2), and default for the remaining settings. Repeat regions identified by RepeatMasker, sites with a mappability score lower than one, annotated sex chromosomes and scaffolds that were not assembled into chromosomes were excluded (Table S5).

#### Global depth filtering

We separated samples passed sample filtering criteria (101 individuals) into two groups according to coverage, referred to as low depth group (16 individuals, coverage < 6X) and as high depth group (85 individuals, coverage > 6X). For each group, we estimated global depth (read count) per site across samples using ANGSD (-minMapQ 25 -minQ 30 -doCounts 1 - doDepth 1 -dumpCounts 1 -maxdepth 5000) and then estimated the per site median depth. Based on the resulting distribution of global depths for any of the two group sets, we excluded sites that had a global depth below 0.5 times the median and above 1.5 times the median from all analyses (Table S5).

#### Excess heterozygosity (HWE) filtering

We filtered out regions with excessive heterozygosity, which are likely caused by reads from paralogous loci being mis-mapped to a single location and other repetitive regions. Using only the Javan banteng samples, we first generated a preliminary file of genotype likelihoods using ANGSD with the GATK model (-GL 2) from common polymorphic sites (MAF ≥ 0.05 and SNP p < 0.000001), base quality at least 30 (-minQ 30), and minimum mapping quality of 25 (- minMapQ 25). Using this genotype likelihood as input to PCAngsd V0.985 (Meisner and Albrechtsen 2018, 2019), we then calculated the per-site inbreeding coefficients (F), ranging from -1 where all samples are heterozygous to 1 where all samples are homozygous, and performed a Hardy-Weinberg equilibrium likelihood ratio test accounting for population structure. Velicer’s minimum average partial test (Velicer 1976) implemented in PCAngsd was used to infer the optimal number of principal components to model the population structure. Based on the per-site inbreeding coefficients for data sets mapped to all three references, we finally discarded windows of 10kb around sites with significant excessive heterozygosity estimates (F < -0.95 and p < 0.000001, Table S5). Downstream population genetic analyses were restricted to the genomic regions that passed all the above-mentioned filters unless stated otherwise.

### Genotype likelihoods (GLs) calculation

For all analyses in which the low-depth samples were analyzed, we used genotype likelihoods (GLs) or single read sampling to account for the genotype uncertainty of calling genotypes (da Fonseca et al. 2016; Garcia-Erill et al. 2022). We estimated GLs by using the GATK model (-GL 2) in ANGSD, inferring the major and minor allele from the genotype likelihoods (-doMajorMinor 1), estimating the allele frequencies (-doMaf 1), and where applicable, calling SNPs using the default likelihood ratio test (-SNP_pval 1e-6) and a minimum allele frequency filter of 0.05 (- minMaf). Additional filtering was performed simultaneously, including only sites that passed the sites filtering, minimum mapping quality of 25 (-minMapQ 25), and minimum base quality score of 30 (-minQ 30).

### Genotype calling, imputation and phasing

We performed genotype calling for datasets mapped to *de novo banteng* assembly, *BosTau9* and *Waterbuffalo* reference genomes using bcftools V1.14 (Li 2011), using samples only passed the sample filtering and genomic regions retained after sites filtering. Based on mapped reads with minimum base quality of 25 and minimum mapping quality of 30, we used the ‘--per-sample-mF’ flag for the pileup, and the ‘--multiallelic-caller’ for the calling. We retained sites after removing both multiallelic sites and indels for downstream analyses.

Due to low depth of some samples, we performed genotype imputation and phasing to remedy genotype missingness and refine the genotypes. To prepare the input, we extracted bi-allelic SNPs from the genotype data mapped to all three references. Imputation and phasing were then performed using BEAGLE v3.3.2 (Browning and Browning 2009) separately for each chromosome. We visualized the distribution of genotype discordance between the original vcf and imputed vcf genotype.

### Population structure and admixture analyses

#### Principal component analysis (PCAngsd)

To investigate population structure, we performed principal component analysis (PCA) for all samples (101 individuals) and only *Bos javanicus* (72 individuals) using PCAngsd (Meisner and Albrechtsen 2018). Beagle format GL files were prepared as input, by applying a minimum minor allele frequency of 0.05.

#### NGSadmix and evalAdmix

With the same Beagle GL file generated from only *B. javanicus* used in PCAngsd, we estimated admixture proportions for each individual using NGSadmix (Skotte et al.2013). We ran NGSadmix with multiple seeds from K=2 to K=8 until convergence, which we defined as a maximum difference of two log-likelihood units between the top three maximum likelihood results. For the converged run of each K value we subsequently calculated the correlations of residuals for each pair of individuals to evaluate the model fit and to test whether the data violated some of these assumptions for K ancestral clusters using evalAdmix (Garcia-Erill and Albrechtsen 2020). We obtained convergence with K from 2-4.

#### Admixture pedigrees of hybrids (apoh)

We further infer the admixture pedigrees of individuals showing signs of recent admixture identified by NGSadmix using apoh (Garcia-Erill et al. 2023). The rationale behind this method is that the descendants in the very first few generations (∼ 5 generations) after an admixture event, will have specific patterns of paired ancestry proportions in their genome, which deviate from the expected under an older admixture by showing an excess of interancestry heterozygosities. After having identified recently admixed individuals, we excluded them from subsequent analyses aimed at inferring historical, or ancient, evolutionary events, e.g. admixture, demographic histories.

### Genetic diversity, population divergence and Runs of homozygosity

#### Heterozygosity

We assessed the genome-wide heterozygosity for all samples. We first estimated the single-individual SFS using realSFS (Nielsen et al. 2012) in ANGSD, and then divided the number of heterozygous sites by the total number of sites for each individual.

#### Global Fst

To infer population divergence, we calculated genome-wide global *Fst* for each pair of populations. We first estimated the likelihood of sample allele frequency (saf files) for each population with ANGSD. For each population pair, we then inferred the pairwise two-dimensional SFS (2d-SFS) from saf files with realSFS and used them as priors for estimating the pairwise Fst using ‘realSFS fst’ in ANGSD, and applying Hudson’s Fst estimator, which is less sensitive to differences in sample size between populations (Bhatia et al. 2013).

#### Runs of homozygosity (ROH)

We estimated runs of homozygosity (ROHs) for the imputed data sets mapped to *de novo banteng* assembly using PLINK (Purcell et al. 2007). We first filtered data sets across all samples with a minimum allele frequency filter of 0.01, with discarding sites of missing call rates exceeding 0.05, and with removing SNP sites that have weird observed heterozygote frequency O(HET) >0.5. We then called ROHs separately for each sample to account for the high variability in depth across samples, keeping only sites for which that sample had data. We ran PLINK allowing at most 1 heterozygous SNP per window within a ROH (--homozyg-window-het 1), and filtered out all variants with missing calls (--geno 0). For validation, we visualized ROHs across the genome and checked visually for signs of long ROHs being broken up. Distinct ROHs were merged within 100 kb distance, and all of resulting ROHs were categorized into five length groups: <= 1 Mb, 1-2 Mb, 2-5 Mb, 5-10 Mb, and > 10 Mb. Two historical banteng (*B. j. javanicus*) samples were not included in this analysis.

### Population history and introgression analysis

#### Treemix

To investigate the history of population splits and historical admixture events, we performed a TreeMix analysis (Pickrell and Pritchard 2012) using the imputed data mapped to waterbuffalo reference to avoid potential internal reference (*de novo banteng* assembly) bias. As relatedness can underestimate covariance and lead to spurious inferences of migration, we applied a more stringent relatedness filtering of a threshold of KING kinship coefficient of 0.1 to exclude potentially second degree of related samples, and further selected five pure samples for each population. Only three samples from each population of historical Javan banteng, Gaur and Africa buffalo were included due to small sample size (n=3). In the TreeMix analysis we included the historical, captive, wild born banteng (*B. j. javanicus*), Bali cattle (*B. j. domesticus*) from Bali, Kupang and Australia, South Asian zebu, European taurine, Gaur, and the Africa buffalo to root the tree. We ran TreeMix assuming 0-1 migration events. For each number of migration events (m), we ran 100 iterations using bootstrap (-bootstrap), a block size of 1000 SNPs (-k 1000), and retained the run with the highest likelihood, breaking ties at random.

#### D-Statistics

To further investigate the presence of ancient gene flow between banteng (*B. Javanicus*) populations and cattle (taurine and zebu), we performed D-statistics (ABBA-BABA) tests (Green et al. 2010; Durand et al. 2011) for data sets mapped to *waterbuffalo* reference to avoid potential internal reference bias, with individual ‘S._caffer_98_608’ from Africa buffalo as the outgroup. Using blocks of a predefined size of 5 MB, we applied the function ‘-doAbbaBaba 1’ in ANGSD between each triplet of H1, H2 and H3 individuals. Block jackknife approach was subsequently used to estimate standard errors and to calculate Z scores (Busing et al.1999). We further performed Bonferroni correction on the inferred raw p-values to take into account the large number of comparisons (Shaffer 1995). Historical samples were not included in this analysis due to relatively high error rates.

#### Outgroup *f*3 statistics

In order to compare the ancient gene flow between historical bantengs and cattle (taurine and zebu) with other banteng populations and cattle, we calculated outgroup *f*3 statistics based on genotype calls using ADMIXTOOLS (Patterson et al. 2012). We first calculated f2 statistics for each population using five million bp blocks (blgsize = 5e6). Using the Africa buffalo as an outgroup, outgroup f3 was estimated in the form of f3 (X, cattles; Africa buffalo), where X represents different populations of bantengs.

### Demographic history

#### PSMC

We estimated the effective population sizes of banteng populations back through time using the Pairwise Sequentially Markovian Coalescent (PSMC) model (Li and Durbin 2011). We ran PSMC on selected 20 pure high depth samples from populations of Captive Javan banteng, Bali cattle from Bali, Kupang and Australia (five samples for each population) with an average depth of 15.2X. In addition to the sites filter, we removed sites based on the average depth per individual divided by three as a minimum and twice the average depth per individual as a maximum. Default settings were used for all PSMC parameters. A mutation rate of 1.26 × 10^−8^ per site per generation and a generation time of 6 years were used to scale the results for visualization (Chen et al. 2018).

#### Fastsimcoal2

To elucidate the domestication history of bantengs (*B. javanicus*), we inferred the joint demographic histories between the wild-type banteng (*B. j. javanicus*) and domestic-type Bali cattle (*B. j. domesticus*) using a coalescent simulation based method implemented in Fastsimcoal2 V2.7.0.9 (Excoffier et al. 2013). The ‘Banteng-WildBorn’ population was used to represent banteng and ‘Bali-Bali’ population was used to represent Bali cattle, based on results from PCA, Treemix and D-statistics, which suggested these populations were least influenced by genetic drift and gene flow. To minimize potential bias arising when determining ancestral allelic states, we used the folded 2d-SFS as input for the inference (Figure S11). Forteen alternative models of historical events were fitted to the joint SFS of Javan banteng and Bali cattle, and we allowed only instantaneous population size changes (Figure S12). For each model we ran 100 independent Fastsimcoal runs to find the best-fitting parameters yielding the highest likelihood, with 500,000 coalescent simulations per likelihood estimation (-n500000), and 100 conditional maximization algorithm cycles (-L100). We then compared the likelihood values obtained under each model and selected the best model based on model normalized relative likelihood (ω_i_), as recommended by Excoffier et al. (2013). To obtain confidence intervals for the parameters, we used a genomic jackknife based on SFS calculated from 100 blocks using winsfs (Rasmussen et al. 2022). For each of those 100, we performed 20 independent Fastsimcoal runs, using the same settings as for the analyses of the original dataset, and calculated the standard error for parameters from the maximum likelihood run from each SFS. The confidence interval for each parameter were defined as ±1.96✕SE from estimates of optimal runs. We used the same mutation rate and generation time as used in PSMC and PopSizeABC for Fastsimcoal2 to convert model estimates from coalescence units to absolute values (i.e. years).

#### SMC^++^

As an independent validation of the divergence time between banteng and Bali cattle, we estimated the divergence time between population of ‘Banteng-WildBorn’ and population of ‘Bali-Bali’ using SMC^++^ (Terhorst et al.2017). SMC^++^ can jointly infer population size histories and split times in diverged populations by employing a novel spline regularization scheme that greatly reduces estimation error (Terhorst et al.2017). Same mutation rate and generation time as used in PSMC and Fastsimcoal2 were used here.

### Signature of selection during domestication

#### Sliding window statistics for *Fst* and π ln-ratio

To identify genomic regions that may have been subject to selection during domestication of banteng, we scanned the genome for regions with extreme divergence in allele frequency (*Fst*) and the highest differences in genetic diversity (π ln-ratio wild/domestic) between banteng (Banteng-WildBorn) and Bali cattle (Bali-Bali) using a genome-wide sliding window approach. Hudson’s *Fst* in 50kbp windows were estimated using the same approach as for global population-pairwise *Fst*. As the selection during domestication will decrease the nucleotide diversity of population, we defined the π ln-ratio as ln(π_Banteng-WildBorn_) - ln(π_Bali-Bali_), where π_Banteng-WildBorn_ and π_Bali-Bali_ are the pairwise nucleotide diversity values for the banteng (*B. j. javanicus*) and Bali cattle (*B. j. domesticus*), respectively. To calculate π ln-ratio, we first generated the likelihood of sample allele frequency (saf files) for each population using ANGSD, and then applied the ‘saf2theta’ function to calculate thetas for each site from the posterior probability of allele frequency. Pairwise nucleotide diversity (π) in 50kbp non-overlapping windows was finally calculated using ‘thetaStat’ in ANGSD and was further used to calculate π ln-ratio. At a significance level of P < 0.005 (Z test, *Fst* > 0.376 and π ln-ratio > 1.33, Figure 5; Table S10), we identified outlier windows as potential selective-sweep regions, used for subsequent analysis and discussion. Any windows that contained < 5,000 sites were excluded.

#### Haplotype based selection scan: XP-EHH

As a complementary approach to identify the selection regions during domestication, we performed genome scan for haplotype based statistics XP-EHH between banteng (Banteng-WildBorn) and Bali cattle (Bali-Bali) using an R package REHH (Gautier and Vitalis 2012). XP-EHH, a cross-population extended haplotype homozygosity, was defined as a method developed to detect selective sweeps in which the selected allele has approached or achieved fixation in one population but remains polymorphic in the other (Sabeti et al. 2007). To search for footprints of selection, SNPs with significantly positive XP-EHH values (XP-EHH > 2.81, Z test, P < 0.005) were considered to be under selection during domestication, and were further checked the overlapping with selective sweep identified by *Fst* and π ln-ratio.

#### Selective sweep regions and their sliding window statistics of Tajima’s D and LD

We defined outlier regions targeted by selection during domestication by overlapping outliers between at least two methods (*Fst*, π ln-ratio and XP-EHH). We then merged these outlier windows into a single region if the distance between windows was < 1Mb. Finally we extracted the genes overlapping with these regions from the de novo banteng reference genome annotation (https://ftp.ncbi.nlm.nih.gov/genomes/all/GCF/032/452/875/GCF_032452875.1_ARS-OSU_banteng_1.0/GCF_032452875.1_ARS-OSU_banteng_1.0_genomic.gff.gz).

To test whether the candidate selective sweep regions identified from *Fst,* π ln-ratio and XP-EHH had an excess of singleton polymorphisms, we computed the Tajima’s D value for both banteng and Bali cattle using the same sliding window approach. Additionally, we also estimated LD represented by the correlation coefficient between pairs of loci (r^2^) using the genotype likelihood based program ngsLD (Fox et al. 2019), with a minimum allele frequency filter of 0.05 and randomly sampling 1% of all SNP pairs for computational feasibility.

### Genetic load

To infer deleterious alleles, we ran SIFT4G (Vaser et al. 2016) on the banteng reference genome assembly (GCF_032452875.1-ARS-OSU_banteng_1.0, see above) and annotated the final VCF file that has been filtered with the resulting SIFT prediction. As SIFT directly sorting tolerant from intolerant protein changes, it gives us more direct estimates of the deleteriousness of the derived allele compared with the reference allele. We used the available GTF file from the banteng reference genome annotation to predict protein, the fasta file of the reference genome, and the UniRef90 fasta file downloaded in January 2024 as protein database source (UniProt Consortium 2023) to run SIFT4G algorithm parallel in 29 chromosomes of banteng genomes. From the available 50,674,907 variants in the filtered imputed VCF file, 2,379,712 variants were assigned SIFT scores. We removed variants that were marked as synonymous but assigned deleterious SIFT scores (<0.05, Vaser et al. 2016) as this may represent sequencing errors or pseudogenes, resulting in a total of 2,377,850 variants with SIFT scores. We limited our analysis on deleterious variants (SIFT <0.05), which are as many as 66,548 variants. This number represents ∼0.1% of the total variants, which is within the expectation of the percentage of deleterious variants in the human genome (Vaser et al. 2016).

Within this set of deleterious variants, we chose variants that are present on all 56 individuals of banteng (Historic, WildBorn, and Captive) and Bali cattle (Bali, Unk-Indonesia, and Australia) to allow comparable magnitude of SIFT score. As the SIFT score is inverted in scale, i.e. the less, the more deleterious it is, we invert it back by making it a subtractor of one so that the more deleterious it is, the higher the score. Then, we multiplied each derived genotype with the SIFT score in each site and summed it for each individual sample. As the maximum score per site would be two from the possibility of two very deleterious alleles in a homozygous derived genotype, the maximum SIFT score would be twice the number of sites, and the SIFT score for each sample can be presented as the percentage of maximum SIFT score.

To determine whether the overall coverage of the samples affect the overall SIFT score of each population, we tested the correlation between the score and the coverage using Pearson’s correlation coefficient in R (cor.test()) (Figure S16 A-B). Due to significant correlation between depth and load score (Figure S16 A-B), we decided to only include samples with coverage larger than 5x and lower than 15x, where load and SIFT scores did not have significant correlations in all types of genotypes inclusion (Figure S16 C-D). After calculating the sum of the total SIFT score per individual, we compare the significance of the different populations only on samples between 5x to 15x depth using one way ANOVA (SIFT score percentage ∼ population) and significant population pairs were emphasised using TukeyHSD() only if they have more than 10 individuals each. All statistical analysis is done in R version 4.4.1. using stats() package.

## Acknowledgements

This work was made possible by and conducted under the permit nr. 210 from year 2024 and extended by the Directorate General of Conservation of Natural Resources and Ecosystem, Ministry of Environment and Forestry of Indonesia. We thank Amal Al-Chaer for significant contribution in extracting DNA samples under challenging conditions. RH, XW and SGA were supported by a European Research Council Starting Grant to RH (Nr. 853442). MHSS was supported by a Carlsberg Foundation Reintegration Fellowship (CF20-0355). We thank Daniel Bradley and M. Thomas P. Gilbert for access to cleanlab facilities, and for assistance in generating the historical genomes. We also thank the Global Species Management Plan Consortium – GSMP Indonesia, and members of the Indonesian Zoos and Aquariums Association (PKBSI) for providing samples.

## Data availability

The raw sequencing data generated for this project have been deposited in FastQ format in SRA with BioProject accession code PRJNA1243309. Banteng assembly generated has been deposited in NCBI with BioProject accession code PRJNA988033. Scripts used to generate all analyses and plots can be found in the github page https://github.com/xiqtcacf/JavaBantengBaliCattles-Scripts. All Supplementary files are available on XXXXXX.

