## SupplementaryFigures for "Population structure and domestication history of the Javan banteng (*Bos javanicus javanicus*)"

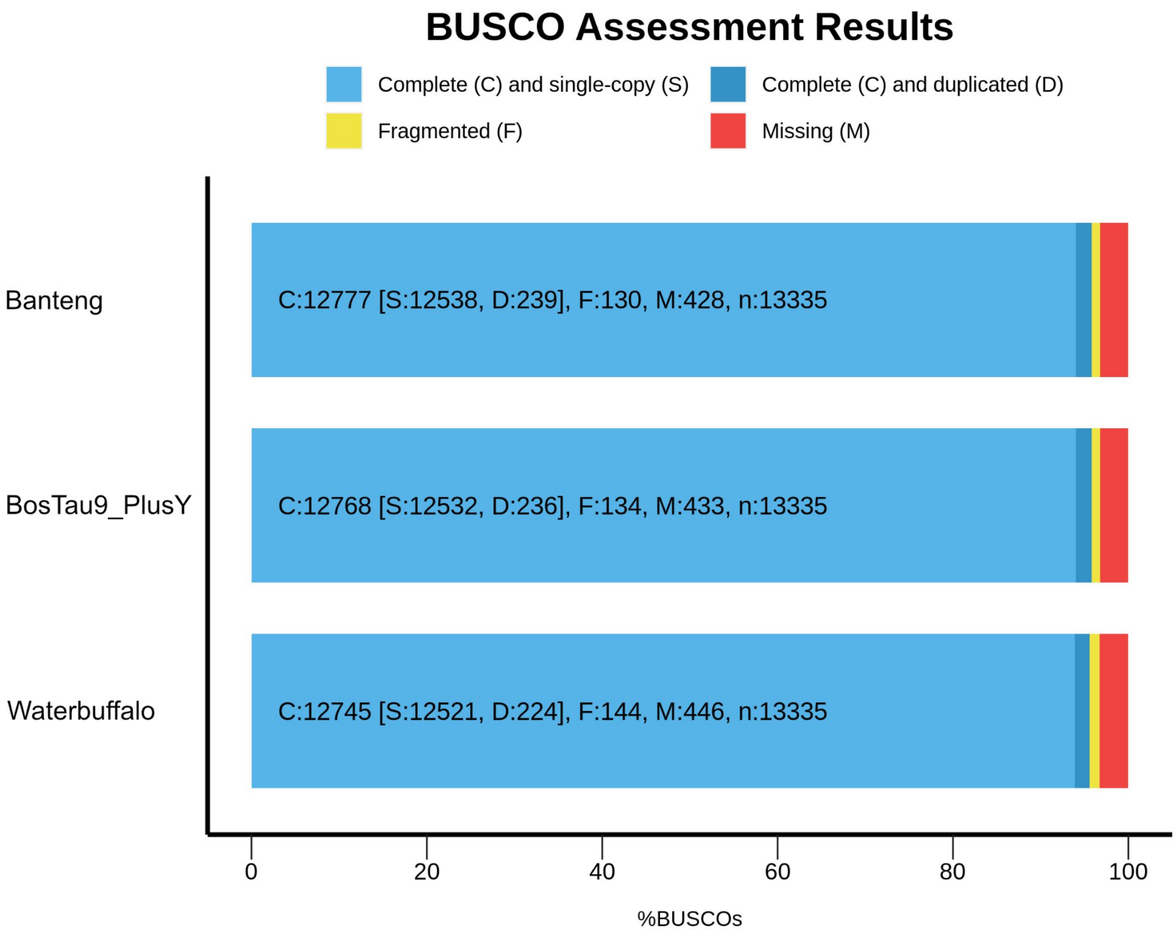


**Figure S1.** BUSCO completeness analysis using lineage dataset ‘cetartiodactyla_odb10’ for three reference used in this study: 1) the *de novo* banteng assembly presented here with MT genome (GenBank: GCF_032452875.1-ARS-OSU_banteng_1.0; MT GenBank: JN632606.1, (2) BosTau9 with Y chromosome (GenBank: GCF_002263795.1-ARS-UCD1.2; Y chromosome GenBank: CM001061.2) and (3) Waterbuffalo (GenBank: GCA_003121395.1-ASM312139v1).

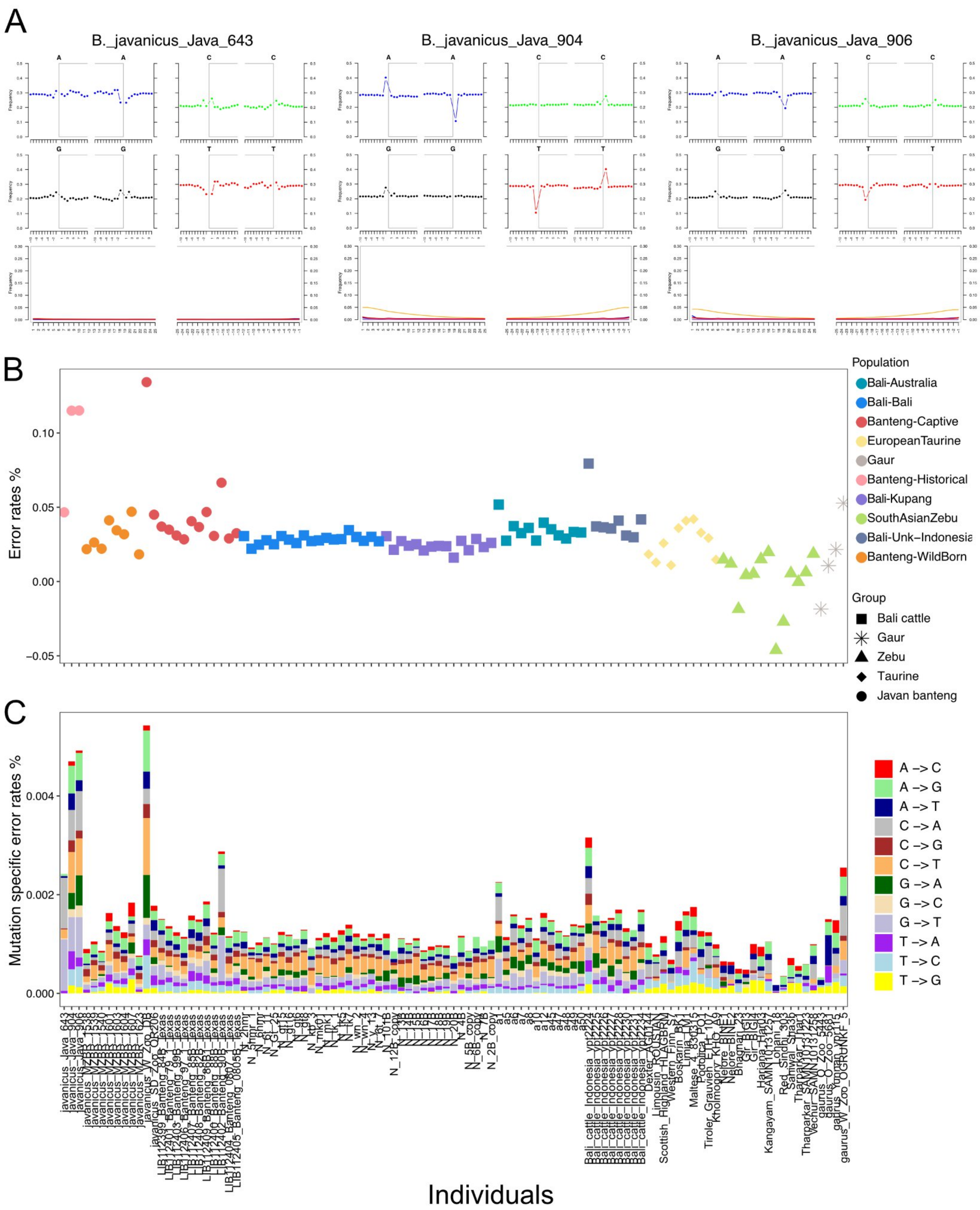
A
B._javanicus_Java_643
B._javanicus_Java_904
B._javanicus_Java_906
B
Error rates %
Population
Bali-Australia
Bali-Bali
Banteng-Captive
EuropeanTaurine
Gaur
Banteng-Historical
Bali-Kupang
SouthAsianZebu
Bali-Unk-Indonesia
Banteng-WildBorn
Group
Bali cattle
Gaur
Zebu
Taurine
Javan banteng
C
Mutation specific error rates %
A -> C
A -> G
A -> T
C -> A
C -> G
C -> T
G -> A
G -> C
G -> T
T -> A
T -> C
T -> G
Individuals

**Figure S2.** Quality assessment for all individuals in this study. (A) Map damage plot for three historical banteng samples. (B) Error rates per individual estimated across all mutations. (C) Error rates per individual estimated by showing mutation specific error rates.

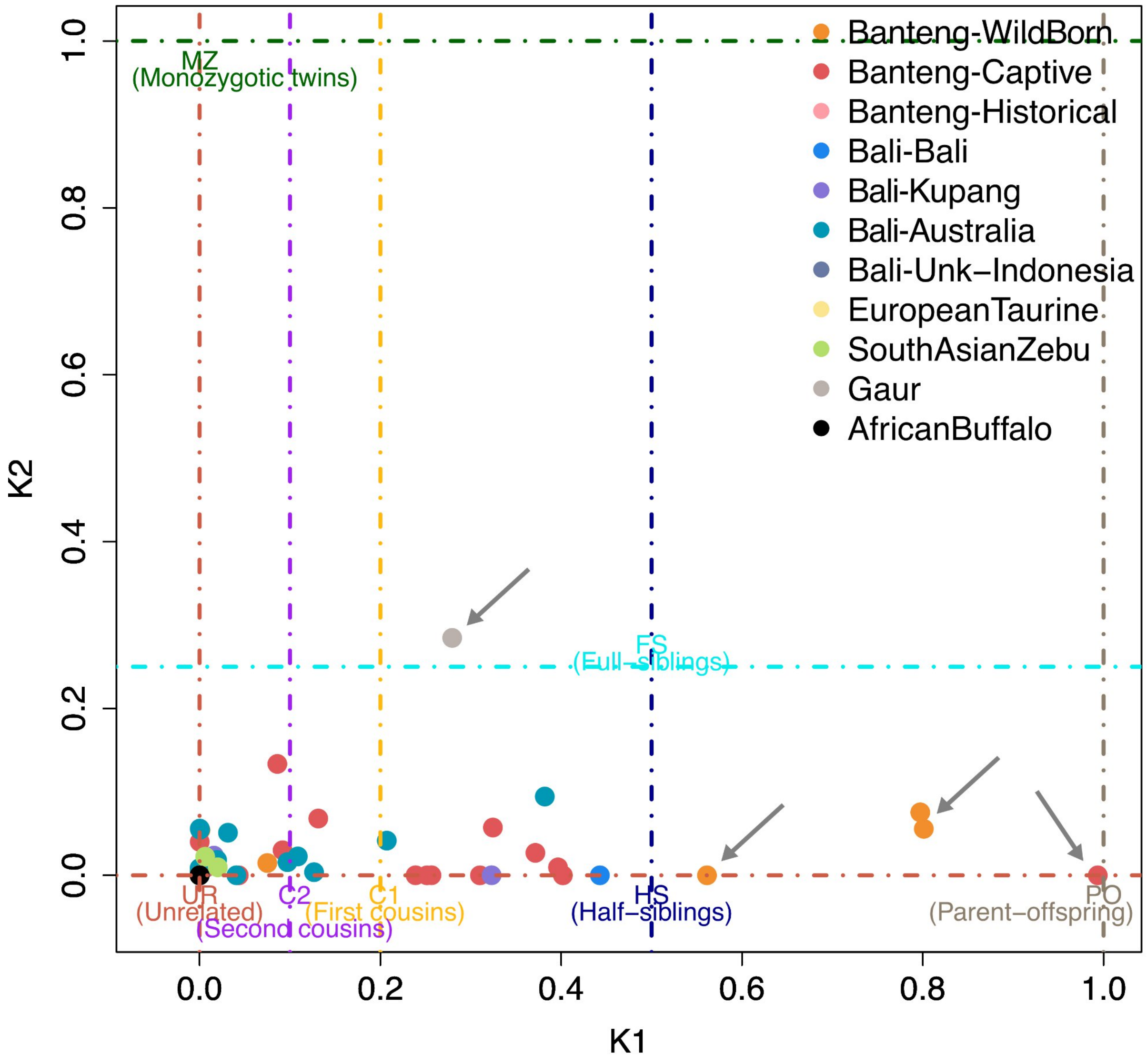


**Figure S3.** Estimates of relatedness for all pairs of individuals within each sampling location or species by NGSrelate. K1 and K2 refers to the proportion of sites where the pair shares one and two alleles identical by descent, respectively.

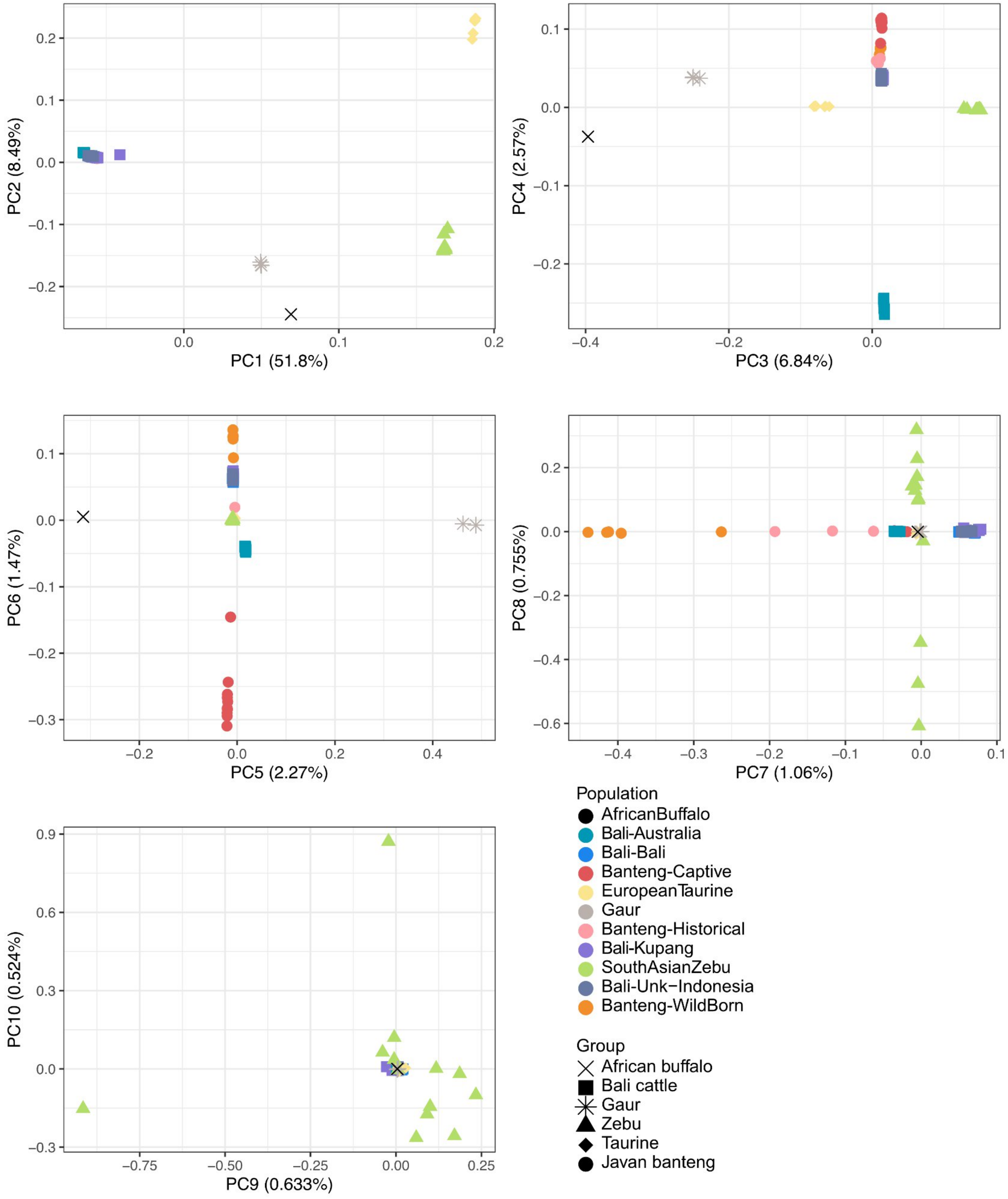


**Figure S4.** PCA plot of all samples colored by sampling locality on the first ten principal components inferred with PCAngsd.

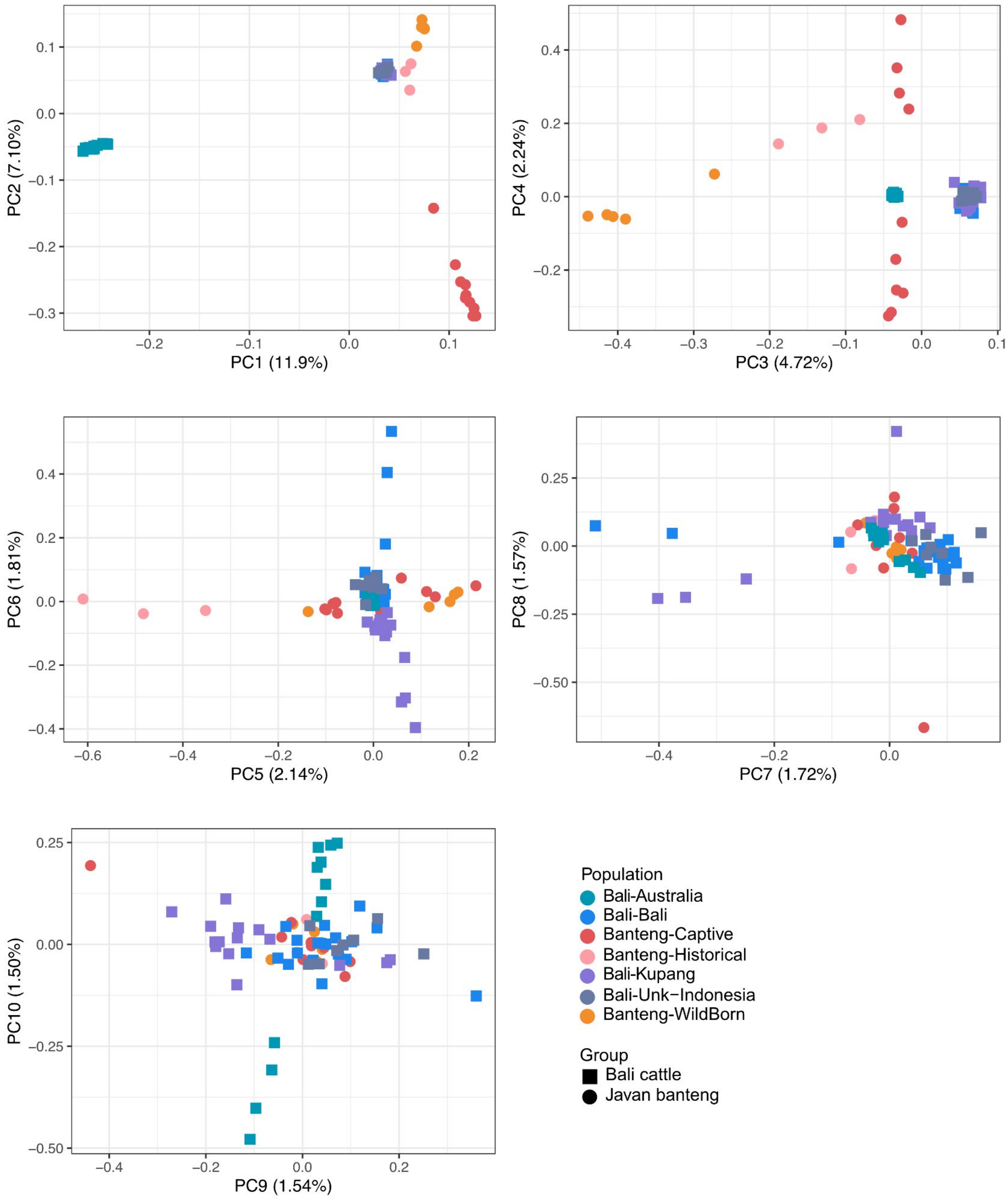


**Figure S5.** PCA plot of banteng samples colored by sampling locality on the first ten principal components inferred with PCAngsd.

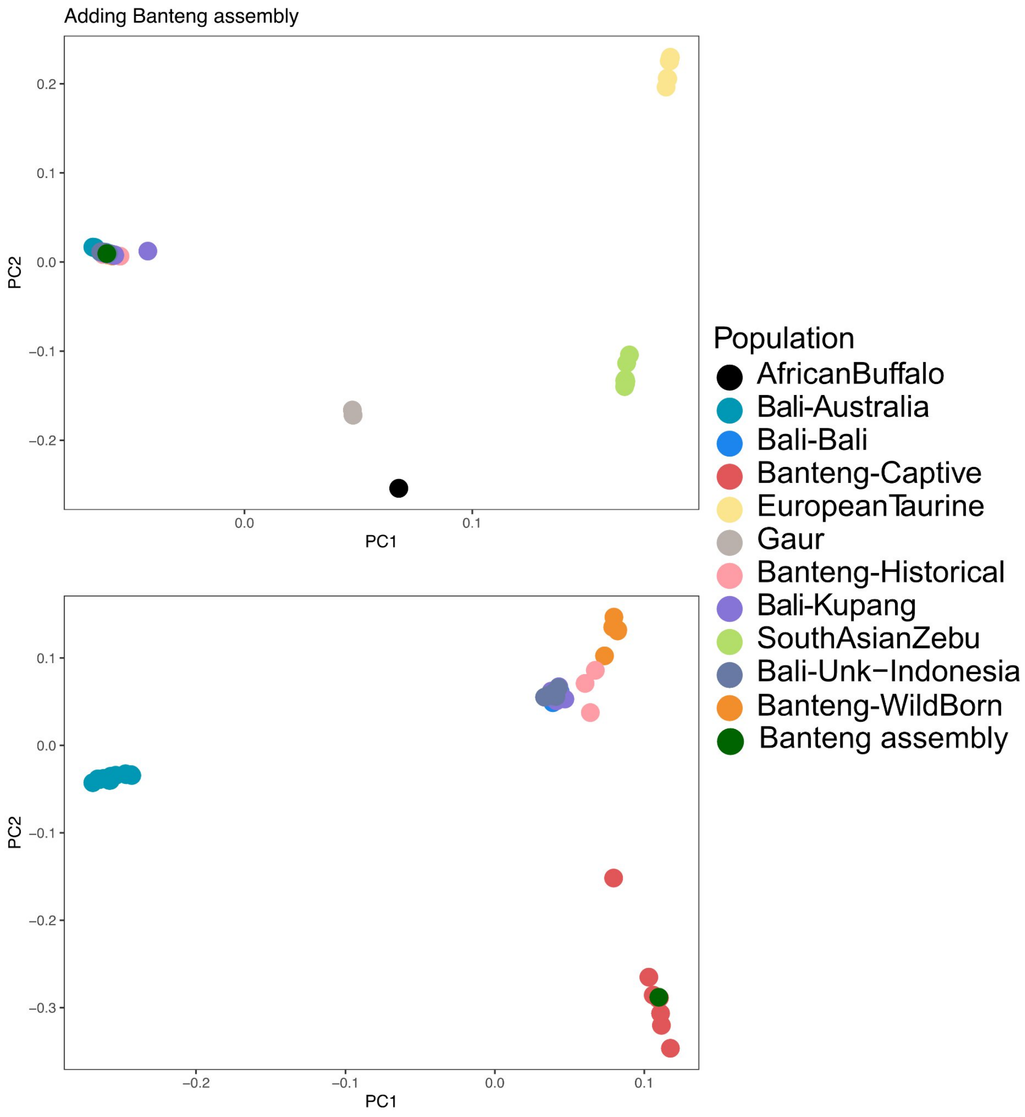


**Figure S6.** PCA plots showing genetic clustering among all individuals (upper panel) and within banteng populations (lower panels) when adding banteng assembly (dark green colour).

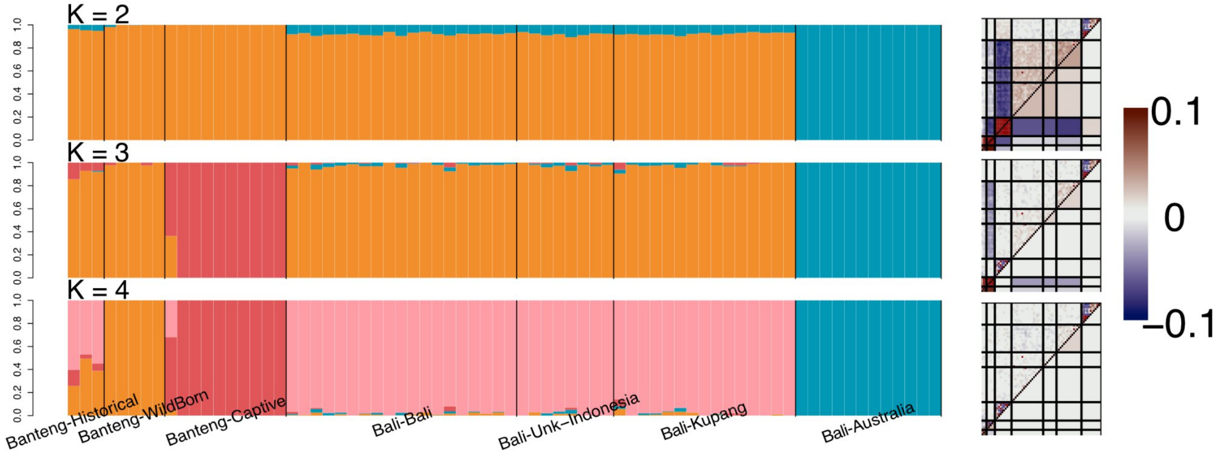


**Figure S7.** Individual admixture proportions estimated with NGSadmix assuming from K = 2 to K = 4. Individuals are grouped by sampling locality. Evaluation of NGSadmix results assuming from K = 2 to K = 4 as the correlation of residuals obtained with evalAdmix.

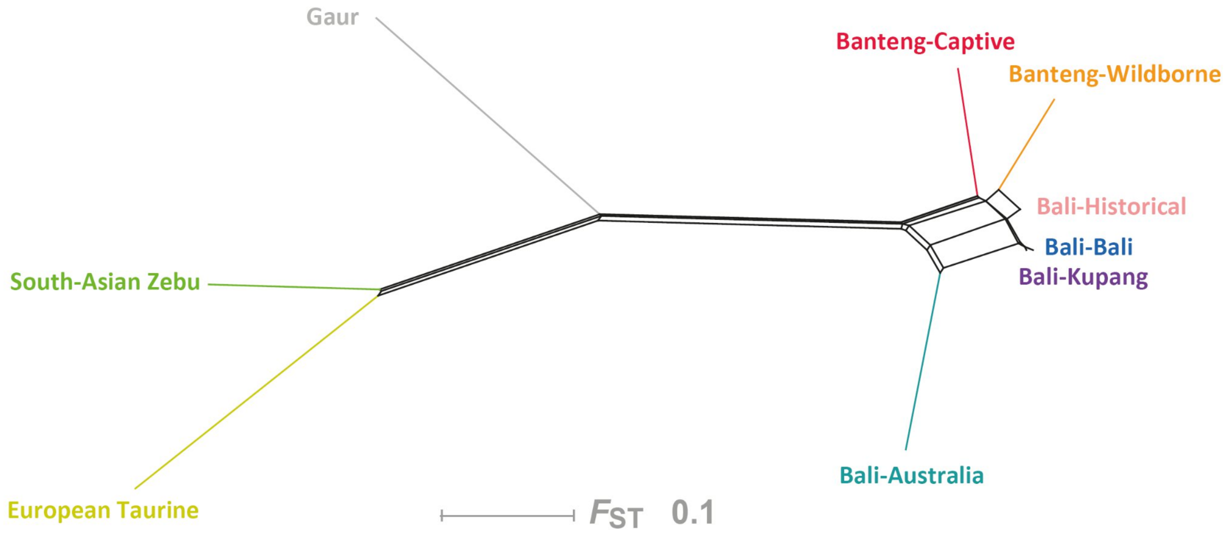


**Figure S8.** Neighbornet visualization of populations based on global Fst values in Figure 2C.

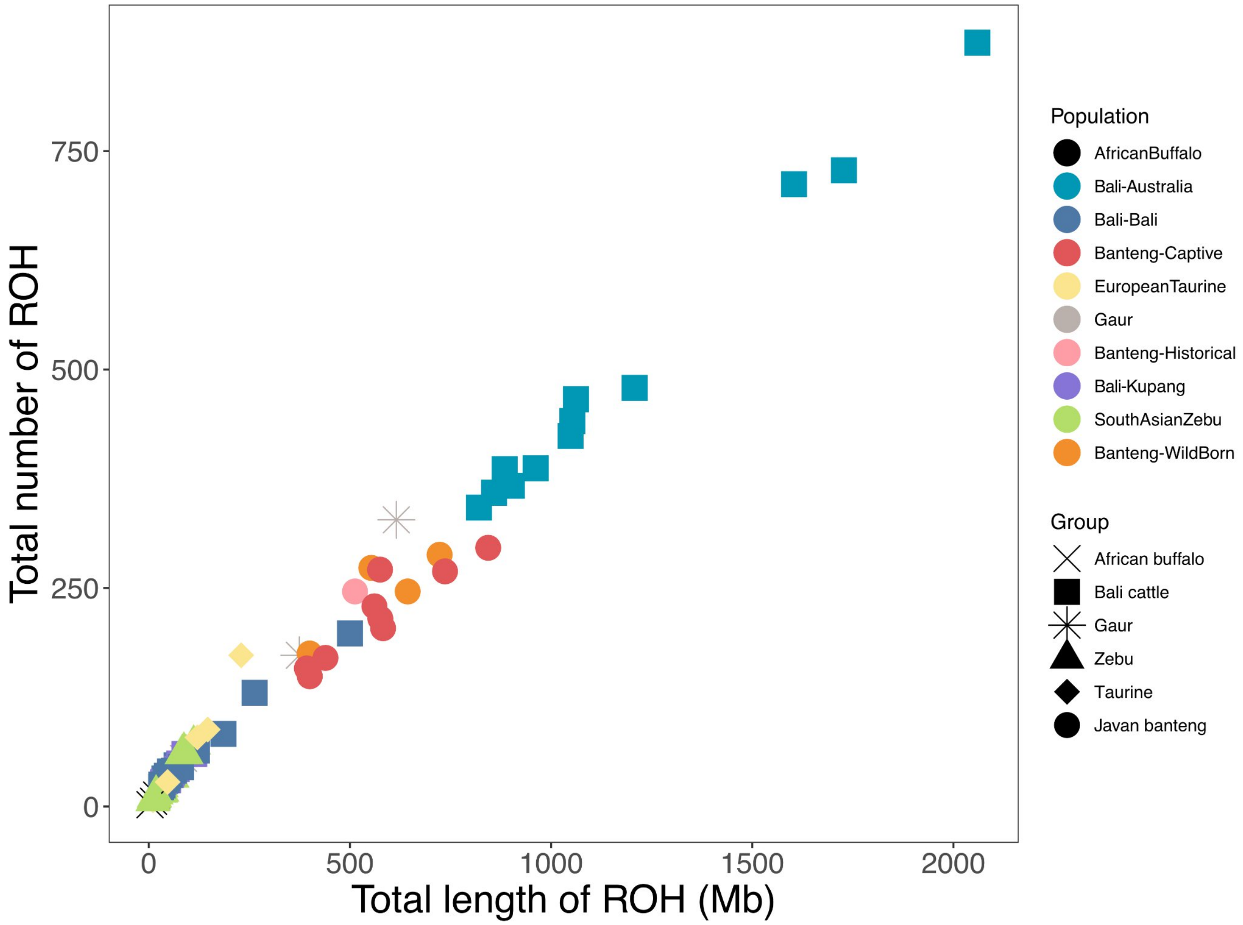


**Figure S9.** The total number of ROH segments (y axis) and the total length (Mb) of the genome in ROH (x axis) for all samples. Each dot represents an individual.

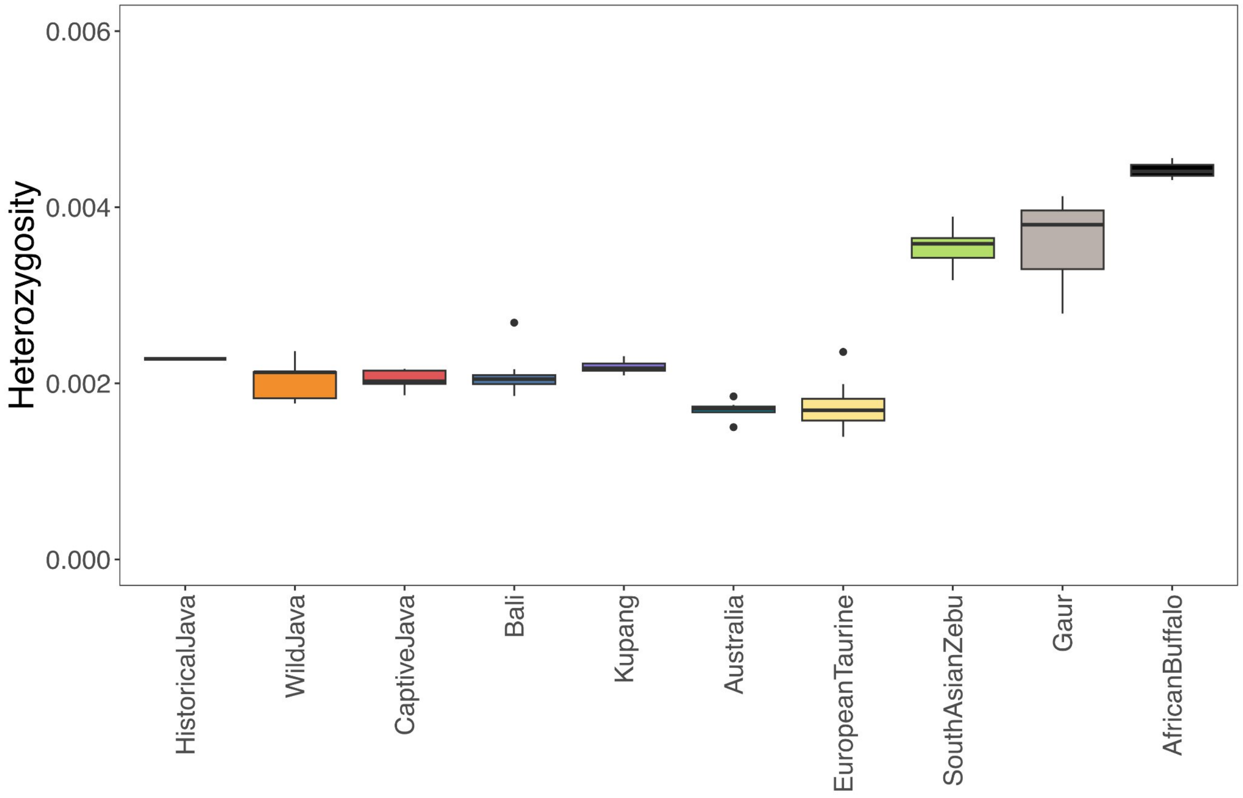


**Figure S10.** Heterozygosity of all individuals excluding ROHs.

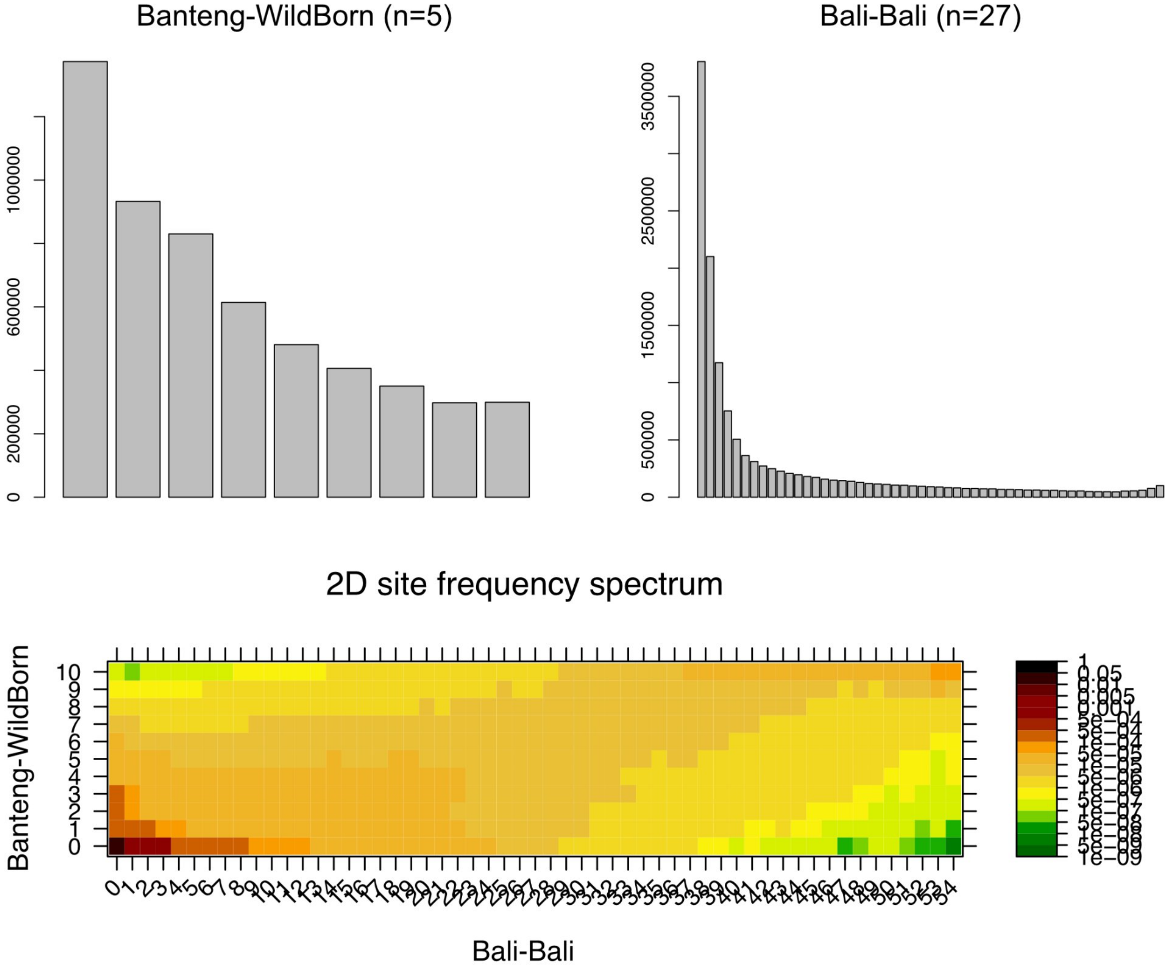


**Figure S11.** 1D-SFS for the population of Banteng-WildBorn and Bali-Bali and the 2D-SFS between two populations.

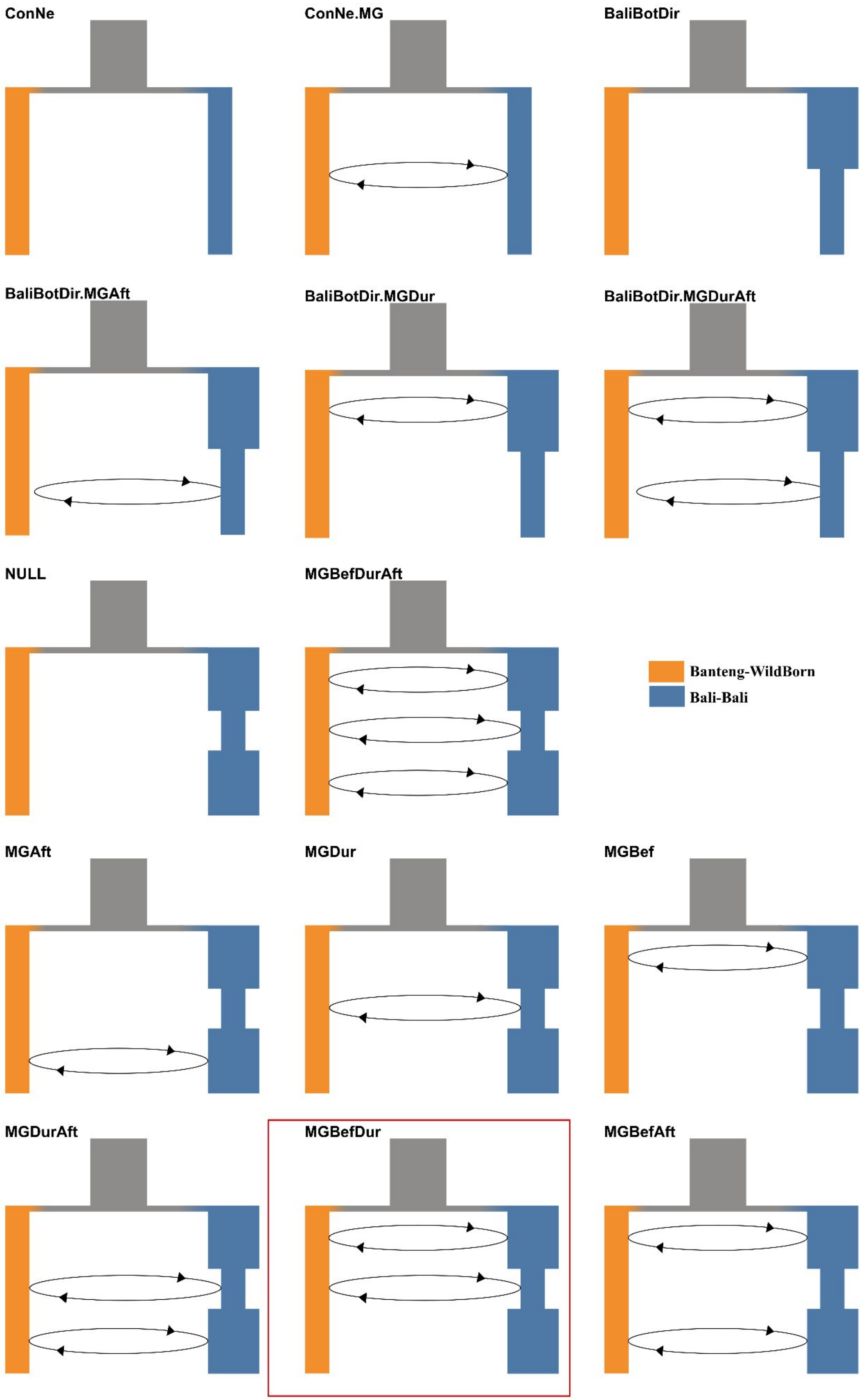
ConNe
ConNe.MG
BaliBotDir
BaliBotDir.MGAft
BaliBotDir.MGDur
BaliBotDir.MGDurAft
NULL
MGBefDurAft
Banteng-WildBorn
Bali-Bali
MGAft
MGDur
MGBef
MGDurAft
MGBefDur
MGBefAft

**Figure S12.** Schematic diagram depicting 14 Fastsimcoal2 demographic models with different assumptions regarding a domestication bottleneck in Bali cattle and the presence of gene flow between Javan banteng and Bali cattle. Model of 'MGBefDur' in red square is the best model based on AIC and model-normalized relative likelihood shown in Table S8.

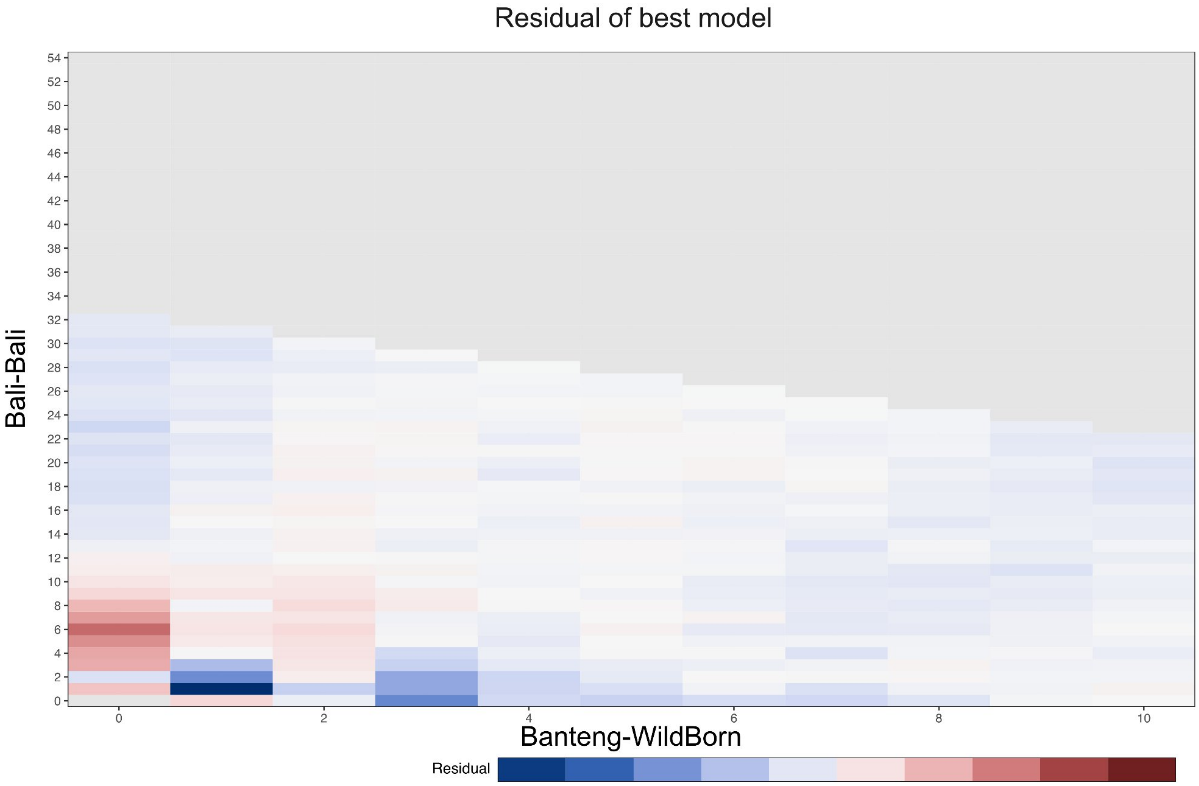


**Figure S13.** Residual between the simulated allele-frequency spectrum under the best model and the observed value from the actual data.

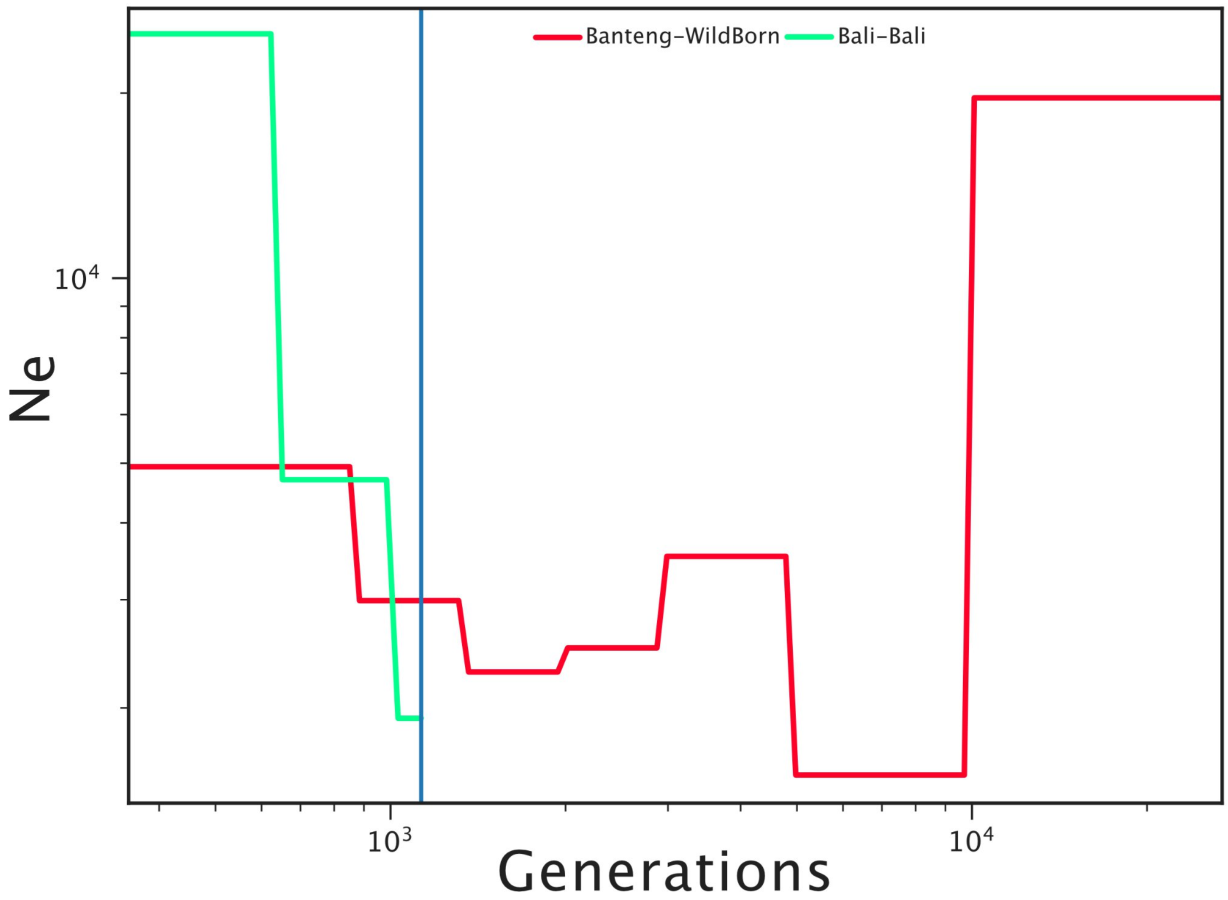


**Figure S14.** Divergence time between wild born Javan banteng (Banteng-WildBorn) and Bali cattle (Bali-Bali), without considering any gene flow between them using SMC++.

**Figure S15.** Tajima's D and linkage disequilibrium (LD) for 49 outlier regions distributed across 19 chromosomes in Table S11.

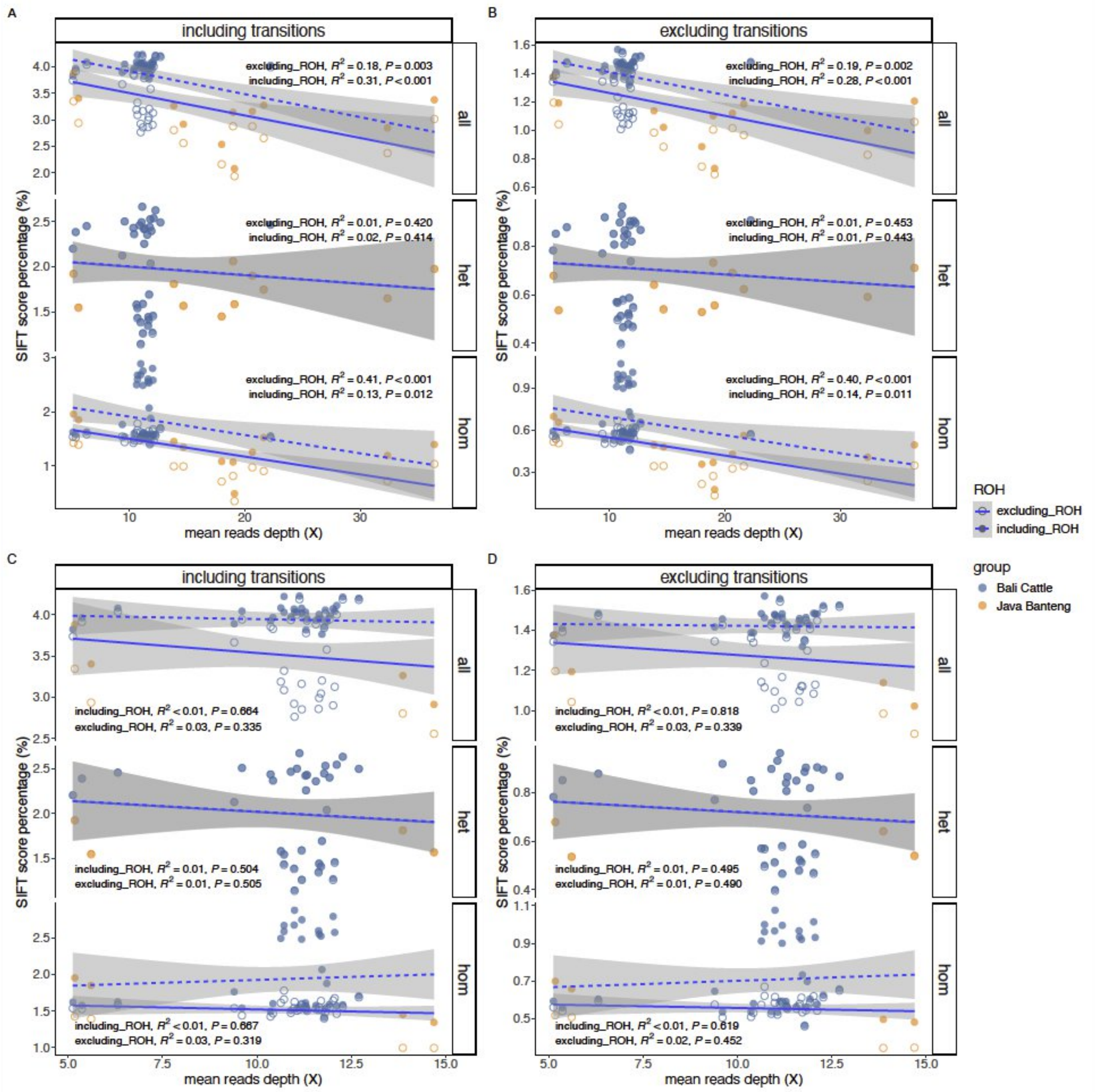


**Figure S16.** Correlation between SIFT scores and mean reads depth (coverage) of all samples (A) while including transitions and (B) excluding transitions to account for damage patterns within the called and imputed genotypes and when limiting to mean reads depth larger than 5x and lower than 15x (C) while including transitions and (D) excluding transitions. Note that SIFT score percentage after excluding transitions decreases ~2 fold.
