## Supplementary figures and images for "Population structure and domestication history of the Javan banteng (*Bos javanicus javanicus*)"

### SupplementaryFigure15

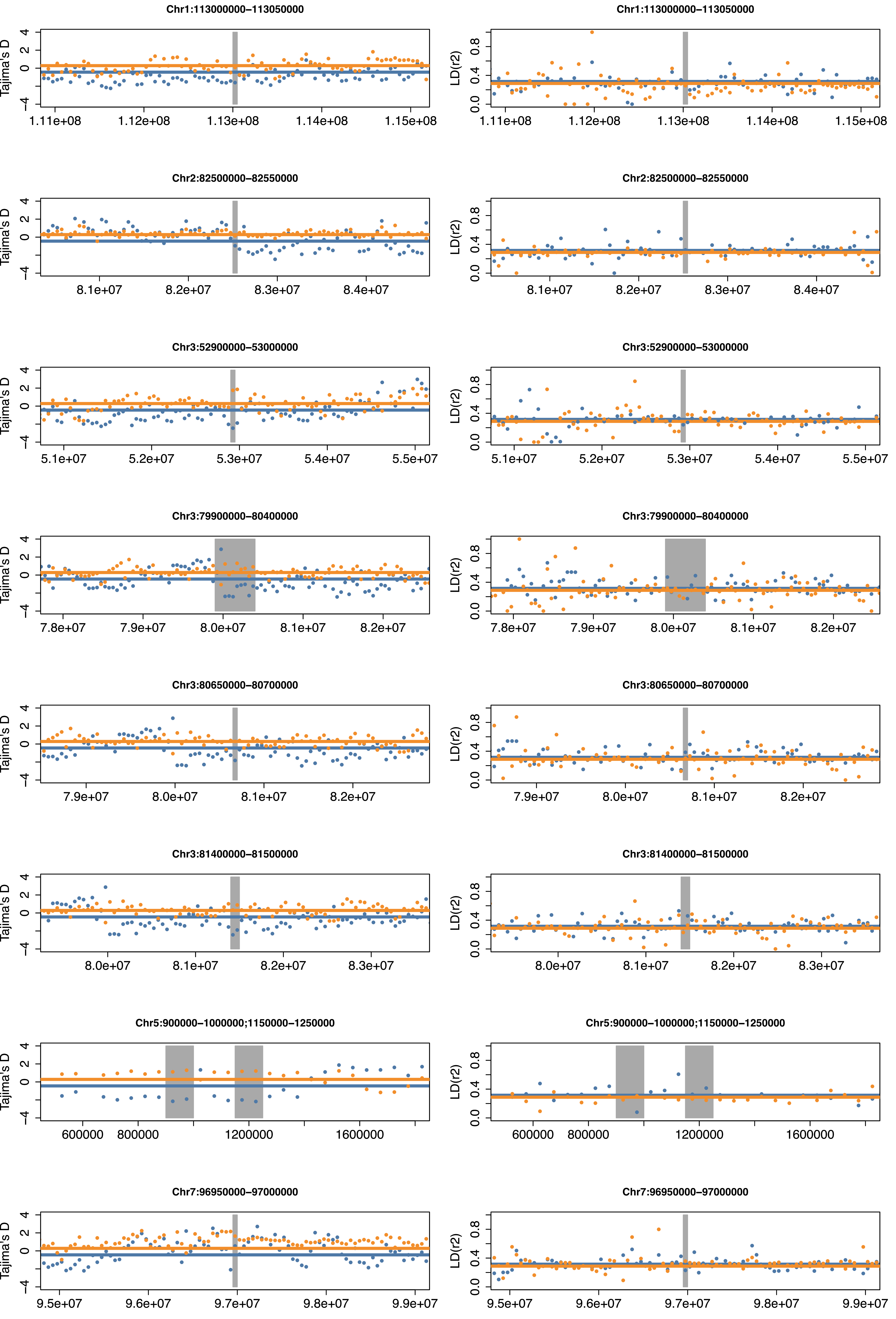

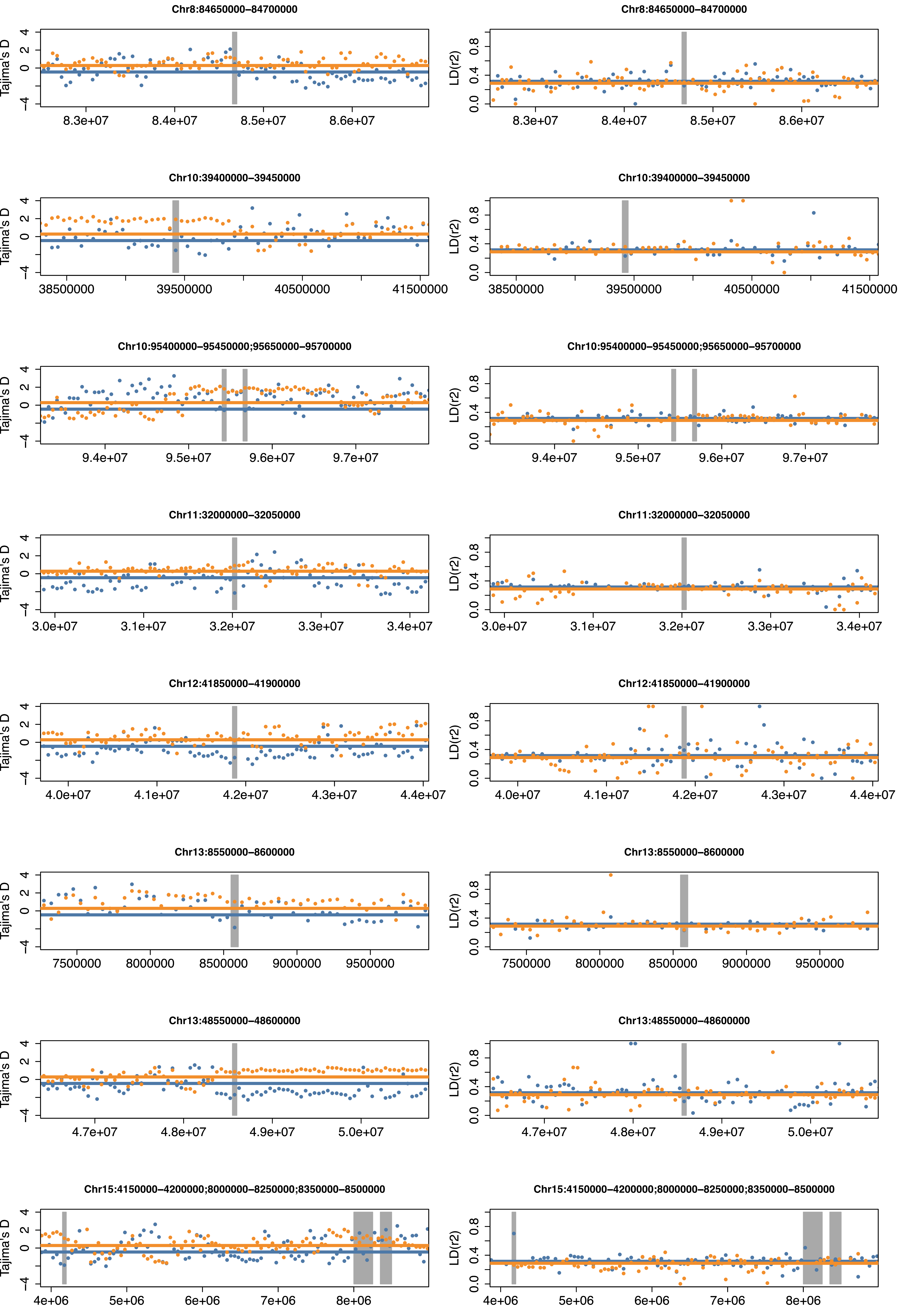

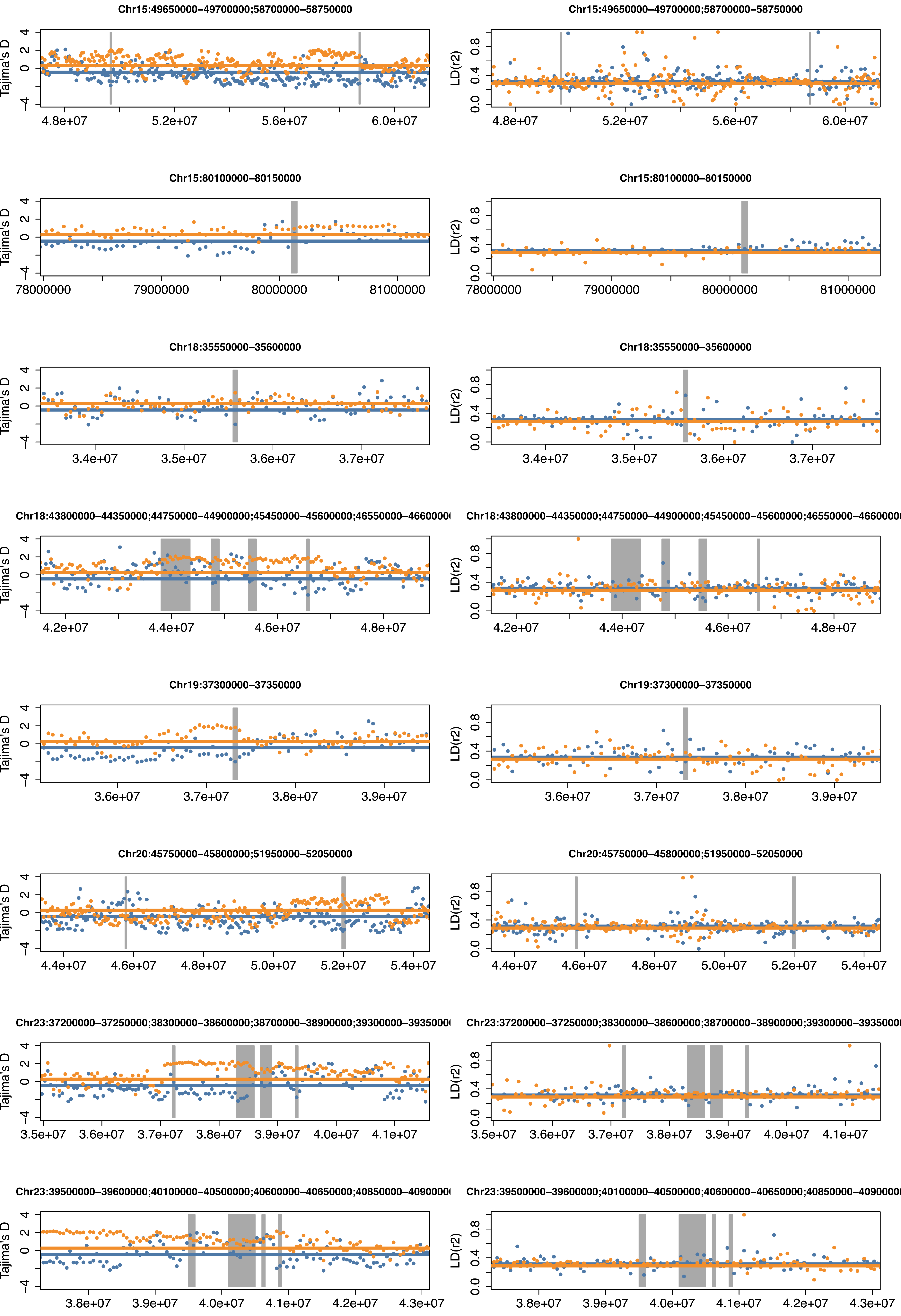

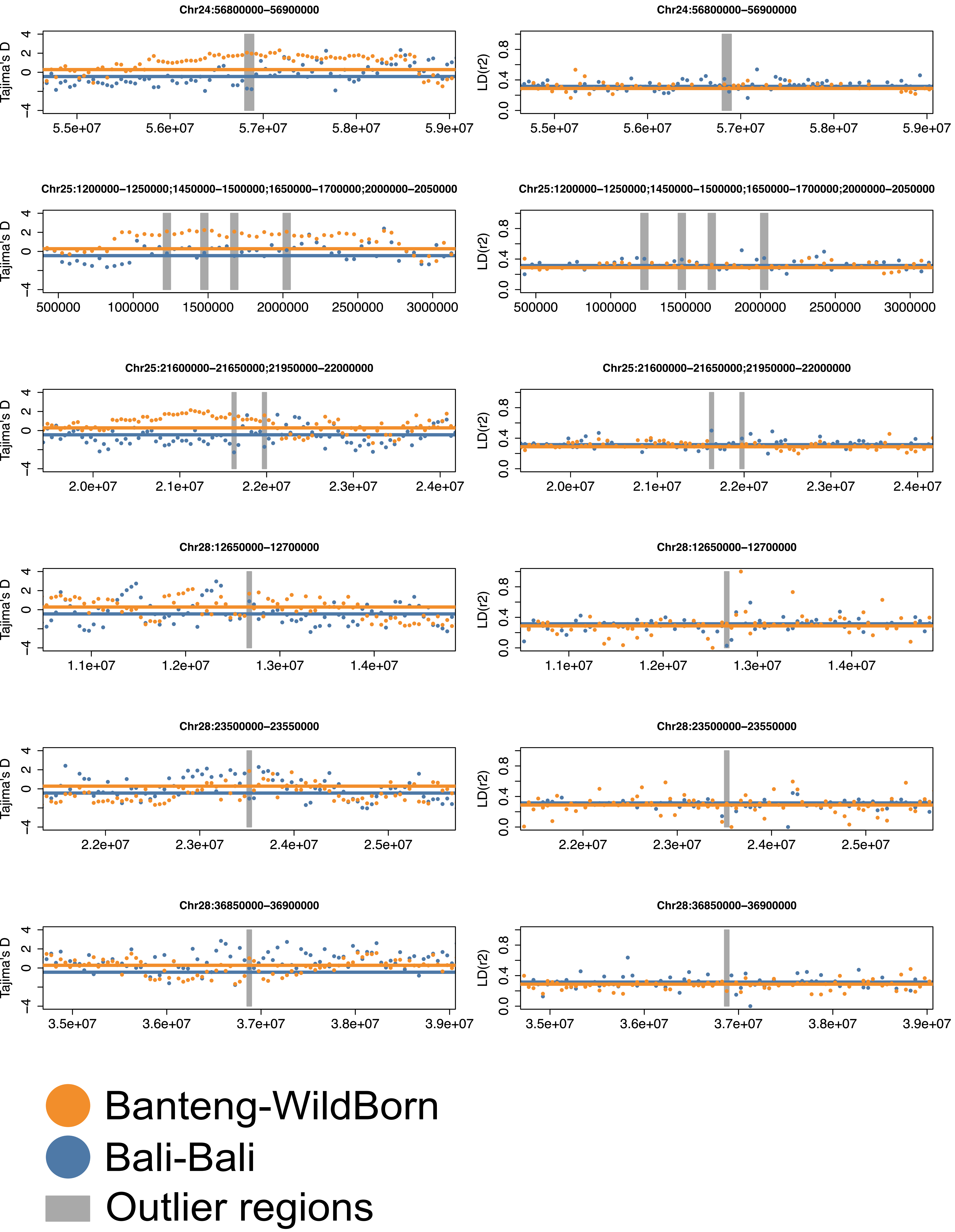
